# FGFR1 inhibition improves therapy efficacy and prevents metabolic adaptation associated with temozolomide resistance in glioblastoma

**DOI:** 10.1101/2025.01.11.632515

**Authors:** Laura Zarzuela, Ignacio G. López-Cepero, Kevin M. Rattigan, Elena Sánchez-Escabias, Macarena Morillo-Huesca, Ana Reina-Bando, Sara S. Oltra, Jesús M. Sierra-Párraga, María Ceballos-Chávez, Vivian Capilla-González, Gema Moreno-Bueno, Piedad del Socorro Murdoch, José C. Reyes, G. Vignir Helgason, Mercedes Tomé, Raúl V. Durán

## Abstract

Recurrent therapy resistance is a major limitation in clinical efficacy and for the outcome of glioblastoma (GBM) patients, positioning GBM among the tumor types with the poorest survival outcomes. In this work, we dissected resistance mechanisms in GBM, which resulted in the identification of FGFR1 pathway as a major regulator of the signaling and metabolic rewiring associated with temozolomide (TMZ) resistance in GBM. Hence, we described a mechanism of resistance that operates at two major levels. First, a p53-mediated regulation of cell cycle inducing cell cycle arrest to allow DNA repair in response to TMZ. And second, a complete metabolic rewiring promoting lipid catabolism and preventing lipid peroxidation. Both the p53-mediated response and the metabolic adaptation are controlled by FGFR1, as inhibition of the FGFR1 pathway completely abolishes this signaling and metabolic reprograming, restoring sensitivity to TMZ. Our results also indicated a correlation of FGFR1 levels with poor prognosis in GBM patients, and validated the treatment of TMZ in combination with FGFR1 inhibitors as an efficient strategy to induce tumor cell death in pre-clinical animal models. This data position the receptor FGFR1 as a very promising candidate for evaluation in future clinical approaches to limit the development of therapy resistance to TMZ in GBM patients.

## INTRODUCTION

Glioblastoma (GBM) is the most aggressive and prevalent malignant primary brain tumor in adults, accounting for 57.7% of gliomas [1]. GBM exhibits one of the worse clinical prognoses, with a median survival of 8-14 months [1], [2]. The current standard of care consists in surgical resection, followed by concomitant radiotherapy and chemotherapy with temozolomide (TMZ), and by adjuvant TMZ treatment [3], [4], [5]. Tumor relapse occurs within 6 months in the majority of GBM patients (90% of total cases), with increased resistance to TMZ and insufficient efficacy of alternative therapies [6], [7], [8]. Indeed, over 50% of GBM patients do not respond to TMZ, either by an inherent or acquired resistance after initial treatment [9], [10]. TMZ introduces methyl groups preferentially at the N-7 or O-6 positions of guanine residues causing DNA damage, replication errors and finally cell death induction [11]. Patients harbouring gene silencing of the enzyme O-6-methylguanine-DNA methyltransferase (MGMT), which reverses TMZ-mediated methylation over the guanine-alkyl groups, partially benefit from TMZ activity [12]. Nevertheless, considering TMZ stability at the acidic pH of the stomach, its complete absortion from the gut, and most importantly, its property to cross the blood-brain barrier [13], TMZ should still be considered as a valuable therapeutic approach, provided that strategies to reverse TMZ resistance are found.

An increasing amount of evidence demonstrate the implication of signaling and metabolic reprogramming in the cellular mechanisms involved in therapy resistance [14],[15], [16], [17]. Particularly, FGFR signaling operates in several compensatory mechanisms driving resistance to EGFR inhibitors (gefitinib, erlotinib) tested in GBM clinical trials [18], [19]. Our previous studies described an FGFR1 and Notch2 signaling crosstalk mediating chemoresistance of neural stem cells (NSC), a likely origin of GBM, suggesting this connection as a potential therapeutic target to overcome therapy resistance [20]. Whether this connection is indeed relevant in GBM resistance to the standard chemotherapy needs to be addressed.

In this study, we have identified that the inhibition of FGFR1 signaling restores GBM sensitivity to TMZ treatment both *in vitro* and *in vivo*. Our results indicated that resistance to TMZ in GBM operated through a p53-mediated program and a metabolic rewiring. FGFR1 inhibition prevented this p53-mediated program, leading to tumor cell death in cell culture and in animal models. Our results also indicated that the synergism of the dual treatment (FGFR1 inhibition plus TMZ) prevented the metabolic adaptation associated with TMZ resistance in GBM. Our analysis confirmed that the success of this novel combinatorial therapy TMZ/FGFR1 inhibitor depends on appropriate GBM patient stratification based on FGFR1 activity and p53 status.

## RESULTS

### FGFR1 and Notch2 activation in human GBM

To investigate the impact of TMZ treatment on FGFR1 and Notch2 signaling pathways we primarily used public cancer genome databases (GEPIA2, http://gepia2.cancer-pku.cn/#index) to analyse the expression of the receptors associated with these pathways and the effect of TMZ treatment in GBM samples. Both FGFR1 and Notch2 showed a significant higher expression in human GBM samples compared to control tissue (Figure 1A), showing Notch2 also an increased expression in low grade gliomas (Supplementary Figure 1A), suggesting a role of FGFR1 and Notch2 specifically in GBM. In addition, none of the other members of the FGFR family (FGFR2, FGFR3 and FGFR4) presented an increased expression in GBM (Supplementary Figure 1B), while Notch1 presented a similar increased expression pattern in GBM as shown by Notch2 (Supplementary Figure 1C). Overall survival analysis showed a worse prognosis for GBM patients with high FGFR1 expression compared to lower FGFR1 expression patients, while no difference in survival was observed comparing low and high expression of Notch2 in GBM patients (Figure 1B), underscoring a direct role of FGFR1 but not Notch2 in GBM aggressiveness. In accordance, a survival heatmap from the genome database analysis showed a significant negative survival correlation only for FGFR1 in GBM (Supplementary Figure 1D). We further confirmed these observations by analysing FGFR1 activation (through FGFR1 phosphorylation) in a set of GBM patient samples by immunohistochemistry (Figure 1C). The analysis of phospho-FGFR1 allowed us to stratify these patients as FGFR1-positive and FGFR1-negative, according to the status of FGFR1 activation in each patient (Figure 1D). A negative correlation was identified between FGFR1 activation and relapse-free survival (p=0.048), with an increased relapse-free survival observed in patients with low levels compared to those with high levels of FGFR1 (Figure 1E). This analysis, therefore, supports a role of FGFR1 activation in GBM aggressiveness.

**Figure 1.**
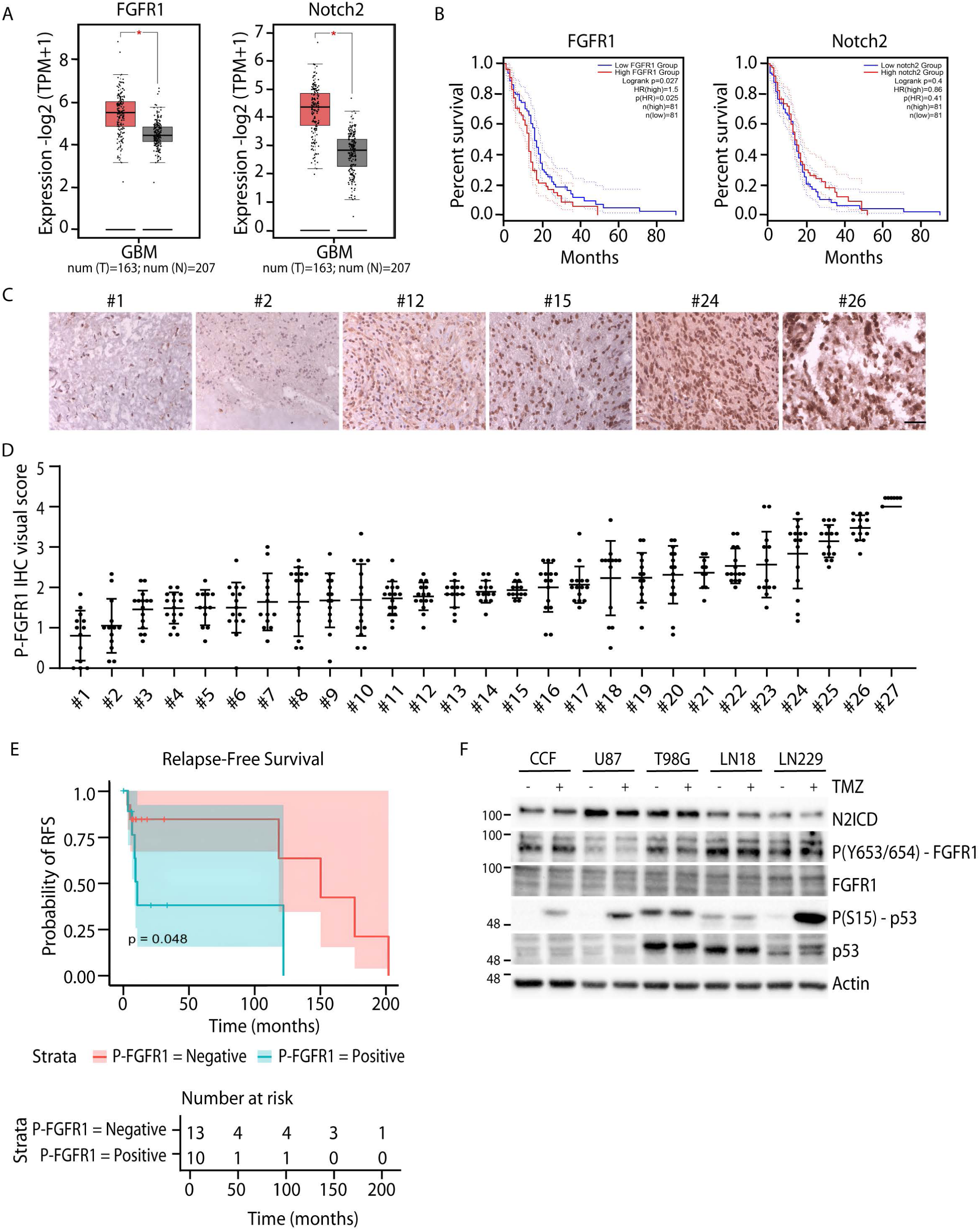
FGFR1 activation correlates with poor prognosis in GBM patients. (A) Box plot representation of FGFR1 and Notch2 gene expression in GBM (T, red) and normal tissue (N, grey) based on tumor and normal samples from the TCGA and the GTEx databases. (B) Kaplan-Meier curve representing overall survival associated with low (blue) and high (red) gene expression of FGFR1 and Notch2 in GBM, obtained as in (A). (C) Representative images of different grades of phospho-FGFR1 expression in human primary GBM samples by immunohistochemistry. (D) Scatter plot for the visual score of images as in (C) for phospho-FGFR1 (P-FGFR1) expression from 27 human primary GBM samples. (E) Kaplan Meier plot curves showing Relapse-Free Survival (RFS) for GBM patients based on the classification as phospho-FGFR1 positive or phospho-FGFR1 negative from the immunohistochemistry samples from (D). (F) Representative immunoblot showing the effect of TMZ treatment on FGFR1 phosphorylation, Notch2 activation (N2ICD) and p53 phosphorylation in several human GBM cell lines.

To further dissect the role of FGFR1 in GBM aggressiveness and as a potential therapeutic target, we analysed a panel of GBM cell lines presenting differential FGFR1 and Notch2 activation (Figure 1F). IC50 analysis in response to TMZ treatment (Supplementary Figure 1E) indicated that the cell lines included in this panel could be classified as high, medium and low TMZ resistant cells (Supplementary Figure 1F). As expected, MGMT+ cells (LN18 and T98G) showed the highest resistance to TMZ, as MGMT is the main resistance mechanism to prevent cell death by TMZ in GBM. The different genetic background of these cell lines (MGMT+ and MGMT- cells; p53 WT and p53 mutant cells; PTEN WT and PTEN mutant cells; Supplementary Figure 1G) did not correlate TMZ resistance to FGFR1 or Notch2 activation, although a tendency was observed for FGFR1 (Supplementary Figure 1H), validating the panel as genetically diverse for comparative purposes. Of note, TMZ treatment did not affect neither FGFR1 nor Notch2 expression or activation (Figure 1F; Supplementary Figure 1I) in GBM cells. We confirmed this result in patient samples using available datasets. FGFR1 and Notch2 gene expression levels obtained from the RNA-seq data of GBM tumors of the TCGA dataset were analysed for patients treated or not with TMZ. TMZ-treated patients did not show increased FGFR1 or Notch2 expression when compared to TMZ-untreated patients (Supplementary Figure 1J). Cell cycle analysis indicated that resistance to TMZ in MGMT negative cells (CCF, LN229 and U87) correlated with cell cycle arrest (Supplementary Figure S1K), and an increase in p53 phosphorylation (Figure 1F). We selected CCF, U87 and LN18 for an in depth analysis to test whether FGFR1 or Notch2 are involved in TMZ resistance in GBM.

### FGFR1 inhibition sensitizes p53 wild-type GBM cells to TMZ

To define the potential role of FGFR1 and Notch2 in TMZ resistance in GBM, we initially assessed the effect of blocking the activity of each pathway, FGFR1 and Notch2, in the sensitization of GBM cells to TMZ. GBM cell lines CCF, U87 and LN18 were treated with either a gamma-secretase inhibitor (DAPT), which blocks Notch receptor activation, or with FGFR1 inhibitors (PD173074 and PD166866). Each type of inhibitor was used either individually or in combination with TMZ for 72h. The combination of DAPT with TMZ did not increase the number of dead cells in any of the cell lines, as estimated by trypan blue assay, suggesting that Notch inhibition was not sufficient to restore TMZ sensitivity (Supplementary Figure 2A-C). To validate the efficacy of DAPT to block Notch cleavage and activation, the levels of both Notch1 intracellular domain (N1ICD) and Notch2 intracellular domain (N2ICD) were analysed by western blot. N1ICD levels were already very low in U87 and LN18 untreated cells, preventing the detection of changes upon DAPT treatment. However, a clear reduction in N1ICD levels was observed in DAPT-treated CCF cells, confirming the efficacy of the inhibitor (Supplementary Figure 2D). Unexpectedly, no effect of DAPT on N2ICD levels was observed in any of the three cell lines (Supplementary Figure 2D), suggesting a lack of efficacy of DAPT to inhibit Notch2 signaling in these cells. Thus, we resorted to Notch2 silencing by siRNA to determine the role of Notch2 signaling in TMZ resistance. Effective Notch2 downregulation using siRNA showed no increase in cell death neither alone or in combination with TMZ (Supplementary Figure 2E-F). These results confirmed a lack of sufficiency of Notch2 to mediate TMZ resistance.

On the other side, the specific pharmacological inhibition of FGFR1 signaling using specific inhibitors (FGFR1i) showed a differential effect among GBM cells. While none of the cells exhibited an increased toxicity by FGFR1i (PD173074) alone, the combination of TMZ and FGFR1i provoked a significant increase in the percentage of dead cells specifically in CCF cells (Figure 2A). On the contrary, the dual treatment using TMZ and FGFR1i did not synergise in cell death induction in U87 and LN18 cells (Figure 2B-C). Flow cytometry analysis of annexin V/PI staining confirmed an induction of apoptosis with >50% of apoptotic cells in CCF cotreated with TMZ and PD173074 while the percentage of apoptotic cells did not significantly increase with the single treatments (Figure 2D-E). The expression of apoptotic markers such as cleaved PARP and cleaved caspase-3 showed an increased expression with the co-treatment specifically in CCF (Figure 2F), further confirming apoptotic cell death in CCF cells treated with TMZ and FGFR1i. Neither of these apoptotic markers were detected with any of the treatments in irresponsive U87 or LN18 cells (Supplementary Figure 2G). Pharmacological inhibition of FGFR1 was confirmed using another specific FGFR1 inhibitor, PD166866. Viability assay and flow cytometry confirmed the ability of PD166866 to increase cell death through apoptosis when combined with TMZ in CCF to a similar extent as PD173074 (Supplementary Figure 2H-I). As observed with PD173074, PD166866 alone or in combination with TMZ was unable to induce cell death in U87 or LN18 (Supplementary Figure 2J). In agreement with the lack of effect of Notch2 inhibition to restore TMZ cytotoxicity, we also observed that, contrary to what we previously described in NSC [20], Notch2 silencing using siRNA did not affect the levels of total and phosphorylated FGFR1 either in the presence or absence of TMZ (Supplementary Figure 2K). Conversely, FGFR1 inhibition downregulated Notch2 signaling, as shown by the decrease in N2ICD levels (Figure 2F), positioning FGFR1 upstream of Notch2 in cell signaling control in GBM cells.

**Figure 2.**
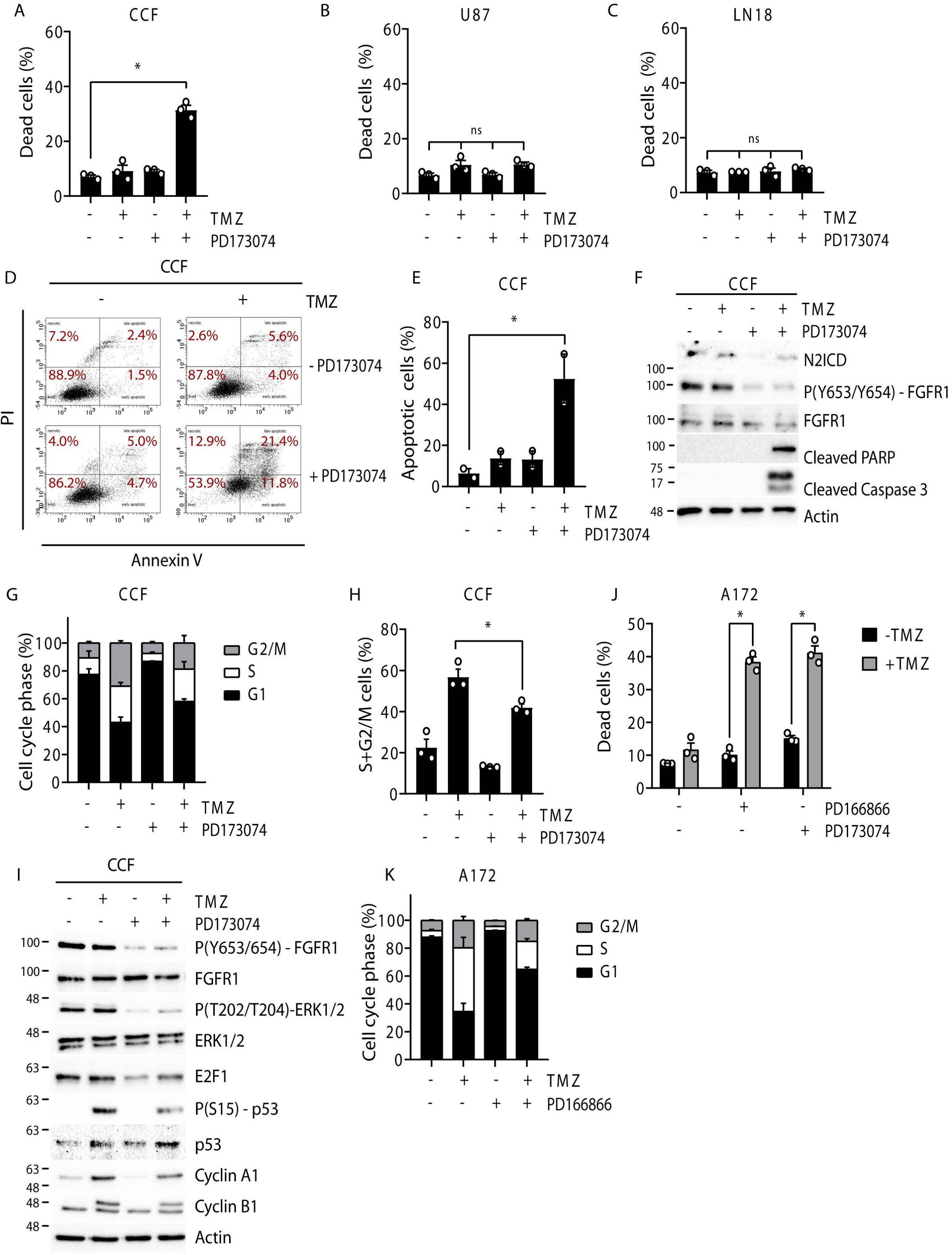
FGFR1 inhibition restores TMZ sensitivity. (A, B, C) Percentage of dead cells as estimated by trypan blue viability assay for GBM cell lines treated with vehicle (DMSO), TMZ (100 µM) or FGFR1i PD173074 (4 µM) for 72 hours. (D) Representative flow cytometry dot plots of annexin V / PI staining of CCF cells treated as indicated for 72 hours. (E) Quantification of the apoptotic population shown in (D) from two biologically independent experiments. (F) Immunoblot of apoptotic markers (cleaved Caspase 3, cleaved PARP), Notch2 activity (N2ICD) and FGFR1 activation (FGFR1 phosphorylation) in CCF cells upon treatment. (G, H) Stacked bar graph (G) of cell cycle analysis by flow cytometry, and quantification (H) of S+G2/M cell percentage for the indicated conditions in CCF cells. (I) Immunoblot of FGFR1 pathway activation (FGFR1 phosphorylation and ERK phosphorylation), and key cell cycle mediators (E2F1, p53 and cyclins) after 72 hours treatment in CCF cells. (J) Percentage of dead cells estimated by trypan blue viability assay for A172 treated with vehicle (DMSO), TMZ (100 µM) and the two FGFR1 inhibitors PD173074 and PD166866 (4 µM and 0.5 µM, respectively), for 72 hours. (K) Stacked bar graph of cell cycle analysis by flow cytometry of A172 treated with PD166866 (0.5 µM) and TMZ (100 µM) for 72 hours. Graphs show mean values ±SEM (n=3 biologically independent experiments). *p < 0.05 (ANOVA post hoc Bonferroni test).

In sight of the cell cycle arrest induced by TMZ previously observed (Supplementary Figure 1K), we next investigated whether the differential capacity of FGFR1i to increase sensitivity to TMZ correlated with cell cycle restoration. Our results indicated that while FGFR1i (PD173074) alone did not affect cell cycle, PD173074 limited TMZ-induced cell cycle arrest (Figure 2G-H). We hypothesized that the FGFR1i-mediated restoration of cell cycle might lead to an increased accumulation of TMZ-induced DNA damage during replication and ultimately to apoptosis. However, DNA damage analysis as measured by γH2AX intensity through automated high content microscopy revealed that the co-treatment FGFR1i + TMZ did not increase γH2AX intensity as compared with TMZ alone, discarding that cell death was due to an increased DNA damage and replication collapse (Supplementary Figure 2L). In view of this result, we further investigated if potential alterations of a key cell cycle mediator such as p53 could be leading to the observed synergistic phenotype. We had previously observed that TMZ-induced cell cycle arrest correlated with an increase in phosphorylated p53 levels in p53 WT cells (Figure 1F). Consistent with a role for p53 in the synergistic phenotype, p53 induction by TMZ treatment was decreased by FGFR1 inhibition using PD173074 or PD166866 (Figure 2I and Supplementary Figure 2M). In agreement with this observation, protein levels of cyclin A1, cyclin B1, cyclin E2 and the p53-related transcription factor E2F1 were also decreased by the co-treatment with FGFR1i, consistent with a p53-mediated cell cycle recovery (Figure 2I and Supplementary Figure 2M). To validate the results observed in CCF cells, we repeated these experiments in A172 cells, another GBM cell line with p53-wild type and FGFR1 positive background (Supplementary Figure 2N). As shown in Figure 2J, similar to what we observed in CCF, co-treatment of TMZ with either of the FGFR1 inhibitors (PD173074 or PD166866) increased cell death, while no effect was observed by the single treatments. TMZ-induced cell cycle arrest, as well as the increased levels of cyclins, were also significantly decreased when co-treated with FGFR1i (Figure 2K and Supplementary Figure 2N). The specificity of FGFR1 inhibition regarding synergism with TMZ with respect to other RTK signaling pathways was confirmed by the lack of effect of the EGFR inhibitor lapatinib. As shown in Supplementary Figure 2O, lapatinib did not mimic FGFR1i and showed no significant effect on cell death upon dual treatment with TMZ. Therefore, the TMZ-mediated resistance program specifically required FGFR1 signalling, positioning FGFR1 as an attractive target to restore sensitivity to TMZ in GBM cells. Similar results were obtained using the ERK inhibitor ravoxertinib (Supplementary Figure 2O), reinforcing the conclusion that FGFR1 is a specific target to restore sensitivity to TMZ in GBM cells. Altogether, these results introduce the concept that p53-wild type GBM restore TMZ chemosensitivity through FGFR1 inhibition.

### FGFR1 controls a transcriptional program involved in TMZ resistance

To gain a better understanding about the molecular events leading to TMZ resistance and further re-sensitization through FGFR1i in GBM cells, we next developed an RNA-seq analysis of TMZ and FGFR1i-treated CCF GBM cells (Figure 3A). Principal component analysis (PCA) of the data highlighted that TMZ treated cells did not differ significantly from untreated cells, while FGFR1i (PD166866) induced a strong change in gene expression pattern in GBM (Supplementary Figure 3A), positioning FGFR1 pathway as a critical regulator of transcriptomic profile in GBM cells. Differential gene expression analysis (unadjusted P value <0.05 and Log2 (FC) >0.5) confirmed that while only 93 genes were identified as deregulated in response to TMZ treatment, the expression of 1180 genes changed in FGFR1i-treated cells compared to untreated cells (Supplementary Figure 3B-C). Dual treatment with TMZ and FGFR1i changed the expression of 1113 genes, with 542 upregulated genes and 571 downregulated genes with respect to untreated cells (Figure 3B). Comparison of FGFR1i and TMZ-treated cells *versus* TMZ-treated cells identified 527 upregulated and 476 downregulated genes (Figure 3C), many of them also found concomitantly deregulated by FGFR1i alone, as shown by the overlapping population in the Venn diagram representation (Supplementary Figure 3D).

**Figure 3.**
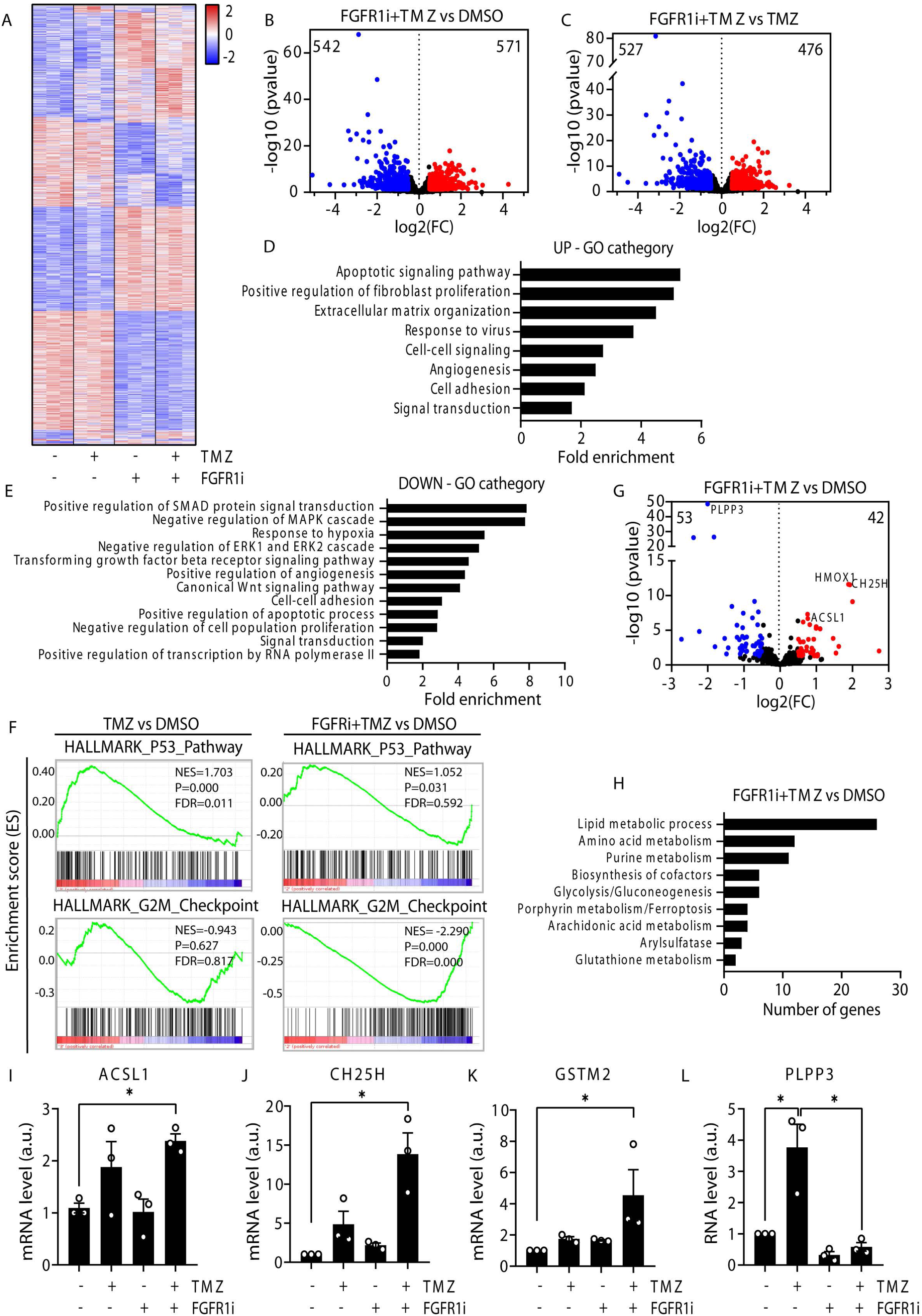
Transcriptomic analysis revealed a metabolic reprograming associated with TMZ resistance. (A) Heatmap of gene expression fold change from RNA-seq analysis in CCF cells treated with TMZ (100 µM) and FGFR1 (PD166866, 5 µM). (B, C) Volcano plot from RNA-seq analysis for upregulated (red) and downregulated (blue) genes for the indicated conditions. (D, E) Gene Ontology (GO) enrichment analysis for biological processes for the up-regulated (D) and down-regulated (E) genes for the dual treatment TMZ plus FGFR1i compared to DMSO (control). (F) GSEA plots for cell cycle related hallmarks comparing gene sets from TMZ treatment and dual treatment with TMZ and FGFR1i. (G) Volcano plot for upregulated (red) and downregulated (blue) metabolism-related genes. (H) GO plot representing the metabolic pathways with dysregulated genes by the treatment with TMZ and FGFR1i. (I-L) mRNA expression levels assessed by qPCR for genes found dysregulated in the RNA-seq analysis and highlighted in (G).

Gene ontology (GO) analysis confirmed that FGFR1i strongly affected MAPK/ERK signaling pathway, but also other signaling pathways such as Smad or Wnt pathways (Figure 3D-E and Supplementary Figure 3E). In support of our previous observations by flow cytometry and immunoblot analysis, gene set enrichment analysis (GSEA) also supported the restoration of cell cycle progression upon the dual treatment TMZ plus FGFR1i as shown by the deregulation of the p53 pathway induced by TMZ and the downregulation of the G2M checkpoint pathway, (Figure 3F). Similarly, the enrichment in upregulated genes related to the E2F1 targets by TMZ treatment, changed to an enrichment in downregulated genes when TMZ was combined with the FGFR1i (Supplementary Figure 3F).

RNA-seq data analysis also revealed that in addition to the expected downregulation of genes involved in cell signaling pathways, FGFR1 inhibition also resulted in a profound change in the profile of metabolism-related genes (Figure 3G-H), with a particular impact in fatty acid metabolism. qPCR analysis confirmed a significant increase in the expression of genes involved in lipid metabolism in cells co-treated with TMZ and FGFR1i, including long chain fatty acyl-CoA synthetase (ACSL1), cholesterol 25-hydrolase (CH25H) or glutathione S transferase Mu2 (GSTM2) (Figure 3I-K). Conversely, in the case of the phospholipid phosphatase 3 (PLPP3), the TMZ- mediated upregulation of this gene was blocked by co-treatment with FGFR1i (Figure 3L). Therefore, our RNA-seq data analysis indicated that the combinatory treatment (TMZ plus FGFR1i) transcriptionally regulated several key pathways affecting cell signalling, cell cycle and metabolism, particularly lipid metabolism.

### FGFR1 inhibition prevents metabolic adaptation associated with TMZ resistance

To validate the potential participation of metabolism in the restoration of TMZ sensitivity upon FGFR1i, we first conducted metabolomics of GBM cells upon TMZ and FGFR1i treatment through LC-MS measurements. TMZ treatment increased the levels of a large number of metabolites including metabolites of the TCA cycle, nucleotide synthesis and amino acid metabolism, while FGFR1 inhibition did not induce any relevant change with respect to untreated cells (Figure 4A). Noteworthy, the levels of glycolytic metabolites were clearly increased in cells treated with both TMZ and FGFR1i as compared to all other conditions (Figure 4A-B). The dual treatment did not significantly change metabolite levels in any other metabolic pathway compared to TMZ alone. Indeed, glycolytic intermediates showed a 2-to-3-fold change increase upon dual treatment FGFR1i plus TMZ (Supplementary Figure 4A), while a slight decrease in the levels of TCA metabolites and amino acids was observed (Figure 4B and Supplementary Figure 4B-C). In accordance with these metabolomics data, glycolysis activity analysed by the extracellular acidification rate (ECAR) showed a further increase in maximal and compensatory glycolysis levels by the dual treatment with respect to TMZ alone (Figure 4C-D and Supplementary Figure 4E). Basal glycolysis also showed a tendency to increase but reaching no significance (Supplementary Figure 4D). Oxygen consumption rate (OCR) analysis showed significant changes in maximal respiration capacity, which was increased in GBM cells in response to TMZ alone and sustained in the dual treatment TMZ plus FGFR1i (Figure 4E-F). A modest increase in basal respiration and mitochondrial ATP production was also observed following the dual treatment although without reaching statistical significance (Supplementary Figure 4F-G). This initial metabolomic and bioenergetic analysis suggested that an increased glycolysis and an increased glycolytic contribution to the bioenergetics of the cell could participate in TMZ resistance and further FGFR1i-mediated re-sensitization.

**Figure 4.**
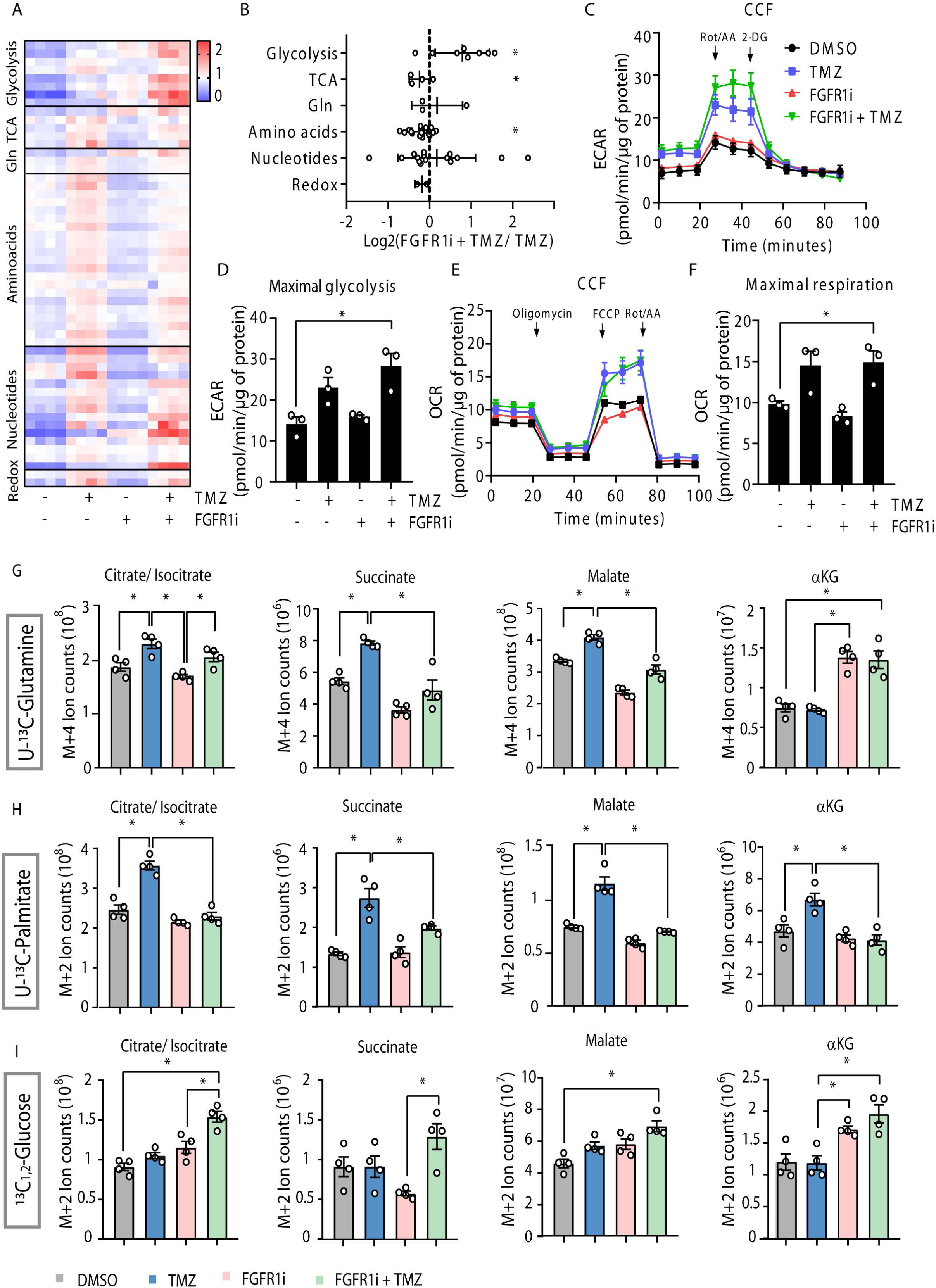
Metabolomic analysis confirmed metabolic rewiring associated with TMZ resistance under the control of FGFR1. (A) Heatmap for metabolite levels, determined by LC-MS analysis, in CCF cells treated for 72 hours with vehicle (DMSO), TMZ (100 µM) and FGFR1i (PD166866, 5 µM). (B) Fold change of the levels of metabolite groups obtained from (A), comparing TMZ and FGFR1i dual treatment with TMZ alone. (C) Representative graph of ECAR analysis measured by Seahorse technology in CCF cells treated with vehicle (DMSO), TMZ (100 µM) and FGFR1i (PD166866, 5 µM). ECAR was measured both in basal conditions and after drug injections (rotenone/antimycin A and 2-DG). Arrows indicate drug injection time. Data show mean values ±SEM (n=3 biologically independent experiments, each performed with 4 or more technical replicates). (D) Maximal glycolysis levels obtained from ECAR analyses, normalized to total protein. (E) Representative graph of OCR analysis measured by Seahorse technology in CCF cells treated as in (C). OCR levels were measured both in basal conditions and after drug injections (oligomycin, FCCP and rotenone/antimycin) to detect basal and maximal respiration capacities. Arrows indicate drug injection time. (F) Maximal respiration capacity levels obtained from OCR analyses, normalized to total protein. (G-I) TCA metabolite isotopologue labelling in CCF cells treated as indicated followed by 4 hours culture with U-^13^C-Glutamine (G), U-^13^C-Palmitate (H) or U-^13^C_1,2_-Glucose (I), determined by LC-MS. M+4/M+5 (G) or M+2 (H, I) ion count levels of the indicated metabolites are plotted. Graphs show mean values ±SEM (n=4 biologically independent experiments). \**p* < 0.05 (ANOVA *post hoc* Bonferroni test).

To validate the possibility of metabolic plasticity as a main contributor to TMZ resistance in GBM, we performed uniformly labelled ^13^C-tracing analysis for the main carbon donors to the TCA cycle, namely glutamine, palmitate and glucose (Supplementary Figure 4H). U-^13^C-glutamine labelling indicated that TMZ increased the incorporation of glutamine to the oxidative TCA as reflected by the increase in labelled (M+4) citrate, succinate and malate (Figure 4G). This TMZ- induced incorporation of glutamine into the TCA was interrupted upon FGFR1 inhibition, as co-treatment with FGFR1i led to an accumulation of M+4 α-ketoglutarate (αKG) but to a decrease of M+4 succinate and malate (Figure 4G). This result, together with the previous metabolomic analysis, strongly suggested that while TMZ promoted the feeding of the TCA cycle from glutamine, FGFR1i prevented the oxidative incorporation of glutaminolysis-derived αKG into the TCA. As previously proposed [21], we hypothesised that this accumulation of labelled αKG and the subsequent decrease in M+4 succinate and malate could reflect an increase in the reductive TCA flux, which therefore would correlate with an increase of the αKG/citrate ratio. Our analysis confirmed that the αKG/citrate ratio was increased upon FGFR1i treatment (Supplementary Figure 4I), further sustaining that FGFR1 inhibition prevents TMZ-induced incorporation of glutamine into the oxidative TCA cycle, and rather forces reductive TCA cycle towards citrate production from αKG. This conclusion was also supported by the observation that the citrate M+5/M+4 ratio was also significantly increased by FGFR1i (Supplementary Figure 4J). Hence, our U-^13^C-glutamine tracing analysis suggested that while TMZ resistance correlated with an increase in the feeding of oxidative TCA cycle from glutamine, concomitant inhibition of FGFR1 reprogramed the fate of glutamine through citrate production via reductive TCA cycle.

Next, to assess the contribution of fatty acid oxidation (FAO) to the TCA cycle in response to TMZ and FGFR1 inhibition, we performed U-^13^C-palmitate labelling metabolomics. As observed in Figure 4H, TMZ treatment increased the contribution of FAO to the TCA cycle as determined by the increase in M+2 labelling of TCA metabolites including succinate, malate and αKG. This TMZ-mediated increase in TCA feeding from FAO was completely abrogated by co-treatment with FGFR1i. Consistent with this observation, the levels of L-carnitine, a shuttle of acyl groups into the mitochondria during FAO, were increased upon TMZ treatment, an increase abolished in cells co-treated with FGFR1i (Supplementary Figure 4K).

In view of the metabolic reprograming observed upon TMZ treatment in GBM, we next aimed at determining whether the TMZ-mediated increase in glutamine and FAO feeding of the TCA was responsible for the chemoresistance. For this purpose, TMZ-treated cells were simultaneously co-treated with glutaminolysis or FAO inhibitors, specifically with the glutaminase inhibitor CB-839 [22] or with etomoxir [23], an inhibitor of CPT1, the enzyme responsible for the formation of acyl carnitines, a limiting step prior to the translocation of acyl groups across the mitochondrial membrane. However, neither CB-839 nor etomoxir treatment were able to induce cell death in TMZ-treated cells (Supplementary Figure 4L). These data suggest that FGFR1i did not restore chemosensitivity to TMZ by just individually blocking one specific metabolic pathway, and rather underscore the role of FGFR1 at modulating a broad metabolic rewiring in response to TMZ.

The fate of glucose carbons was also analysed using ^13^C_1,2_-glucose. The analysis of the labelled carbons showed that TMZ treatment did not alter the contribution of glucose to TCA. However, in agreement with the results obtained above, combinatorial treatment with TMZ and FGFR1i led to an increase in the TCA intermediates derived from glucose (Figure 4I). This result together with the increase in glycolysis observed by metabolomics, suggested a model in which TCA feeding from glucose metabolism compensated for the decreased contribution of both glutamine and fatty acids upon FGFR1 inhibition in TMZ-treated cells, correlating with increased chemosensitivity. Our metabolomics study indicates that FGFR1i did not generate a metabolic collapse in the GBM cells upon co-treatment with TMZ, but rather induces a blockade in the metabolic adaptation associated with TMZ resistance, ultimately leading to cell death.

### FGFR1i-mediated chemosensitization to TMZ affected lipid profile and increased lipid peroxidation in GBM

Based on our metabolomic analysis, we next investigated the implication of a potential lipid metabolism imbalance or reprograming in chemosensitization in GBM. Our transcriptomic data already showed changes in the expression of genes involved in lipid metabolism in response to FGFR1i in TMZ-treated cells (Figure 5A). Thus, we tested the possibility that cell death was accompanied by an imbalance in lipid composition in this condition. First, we labelled GMB cells with the lipid probe Bodipy FL C12 to measure esterification and lipid droplet content. Esterification and lipid droplet formation is a way to control free fatty acid levels, aimed at preventing a dangerous oxidative state. TMZ treatment showed an increased incorporation of this probe compared to untreated cells, and this increase was not blocked by co-treatment with FGFR1i (Figure 5B). This result supports a role of fatty acid uptake, esterification and lipid droplet formation in cell adaptation to TMZ that is not compromised by FGFR1 inhibition. As choline and its derivatives are structural constituents of the cellular membrane, regulating its integrity [24], we next identified alterations in the levels of choline derivatives. Treatment with TMZ significantly increased choline and its derivatives levels (Figure 5C), reinforcing the idea that TMZ resistance operated through a lipid rewiring in GBM cells. Sustaining this possibility, re- sensitization to TMZ by FGFR1i came along with the observation that the inhibition of FGFR1 prevented this TMZ-induced increase in choline derivatives, particularly in the case of choline and phosphocholine (Figure 5D). We therefore specifically explored cell lipid composition under the different conditions by untargeted lipidomics to identify potential differences responsible for the cell death in the combined treatment (Figure 5E). The most striking change was the significant increase in relevant polyunsaturated fatty acids (PUFAs) in the combined treatment compared to all the other conditions, including arachidonic acid (20:4), eicosapentaenoic acid (20:5), docosapentaenoic acid (22:5), adrenic acid (22:4), docosahexaenoic acid (22:6) and nervonic acid (24:1) (Figure 5F-J). It is well known that PUFAs are prone to lipid peroxidation, as oxidants could attack C:C double bond containing lipids [25]. The staining with the lipid peroxidation sensor Bodipy 581/591 C11 showed higher levels of oxidised lipid species in GBM cells (both CCF and A172) treated with TMZ and FGFR1i with respect to any other conditions (Figure 5K-L and Supplementary Figure 5A-B). These results propose that FGFR1i-mediated re- sensitization to TMZ was accompanied by an increase in PUFA peroxidation, which is generally related to ferroptosis [26], [27]. This conclusion was further supported by our transcriptomic analysis, which indicated that FGFR1i plus TMZ treatment induced a significant increase in a number of genes related to ferroptosis, including HMOX1, NOX4 and ACSL1 (Figure 5M), suggesting the implication of ferroptosis in cell death associated with re-sensitization to TMZ by FGFR1 inhibition. In view of these last observations suggesting the participation of ferroptosis in FGFR1i-induced chemosensitization, we further investigated the mechanism of cell death triggered by the cotreatment of TMZ and FGFR1i. GBM cells cotreated with TMZ and FGFR1i were additionally incubated either in the presence or the absence of the pan-caspase inhibitor Q-VD-OPh (QVD) or the ferroptosis inhibitor ferrostatin-1 (Fer-1) [28]. Interestingly, only the combination of both the caspase and the ferroptosis inhibitors was able to abrogate cell death induced by FGFR1i and TMZ co-treatment (Figure 5N and Supplementary Figure 5C). This result further confirmed the participation of lipid peroxidation and ferroptosis, together with apoptosis, in the re-sensitisation of GBM cells to TMZ upon FGFR1 inhibition, highlighting the broad role of FGFR1 in both signaling and metabolic adaptation during TMZ resistance.

**Figure 5.**
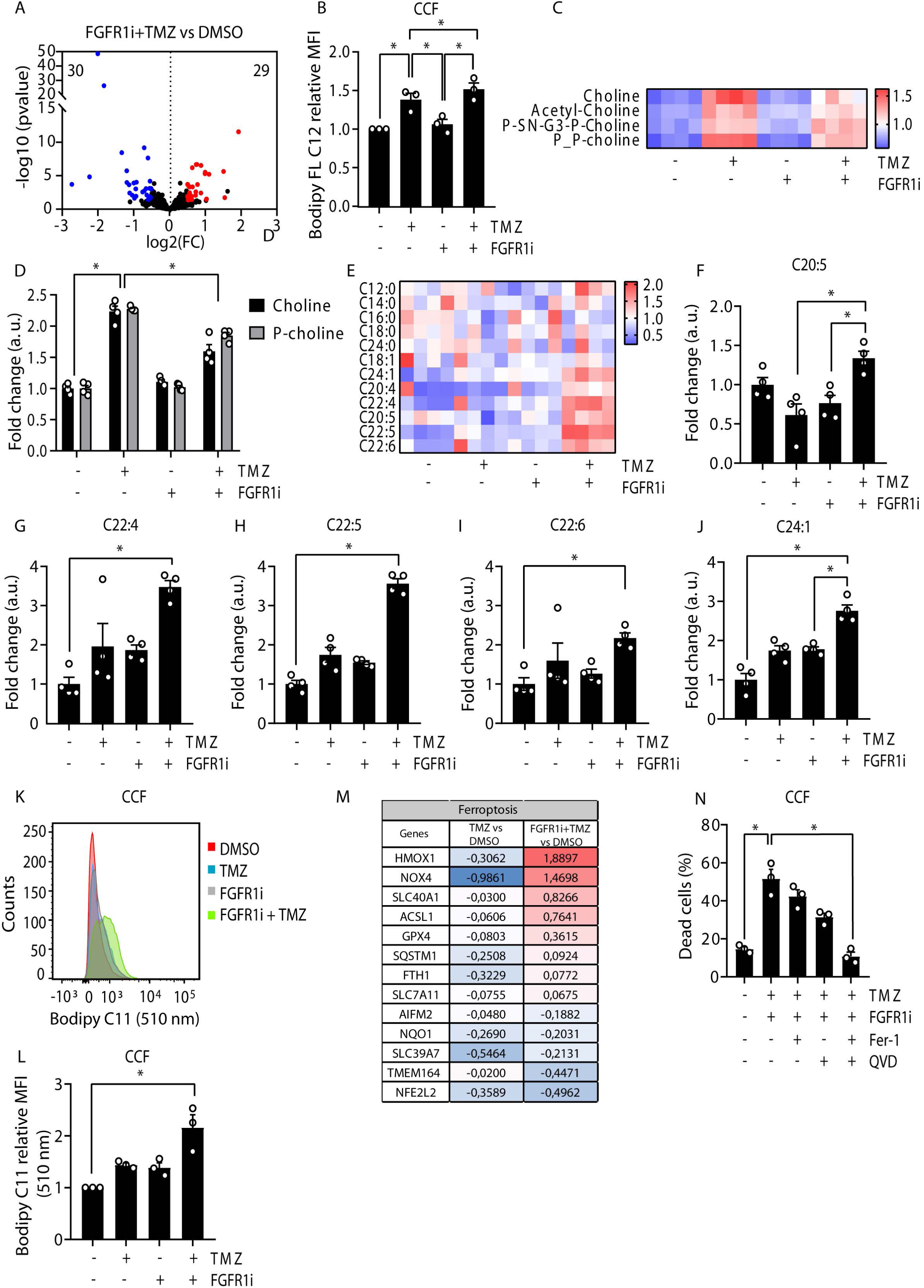
Lipidomic analysis identified lipid profile changes associated with TMZ sensitivity. (A) Volcano plot of lipid metabolism related genes differentially upregulated (red) and downregulated (blue) in the TMZ and FGFR1i cotreatment obtained from RNA-seq analysis in CCF. (B) Lipid droplet quantification by relative mean fluorescence intensity (MFI, 510 nm) using Bodipy FL C12 (2 µM) in CCF cells treated with vehicle (DMSO), TMZ (100 µM) and FGFR1i (PD166866, 5 µM) for 72 hours. (C) Heatmap for choline and derivatives levels determined by LC-MS analysis, in CCF cells treated as indicated for 72 hours. (D) Fold change of choline and phospho-choline (P-choline) for the different treatments obtained from the lipidomics in CCF cells. (E) Heatmap for relative changes of free fatty acids determined by lipidomic analysis in CCF cells treated with vehicle (DMSO), TMZ (100 µM) and FGFR1i (PD166866, 5 µM) for 72 hours. (F-J) Bar plot representations of each indicated unsaturated fatty acid as obtained for lipidomic analysis. (K, L) Representative flow cytometry histogram (K) and mean fluorescence intensity quantification (L) showing lipid peroxidation levels as determined by Bodipy 581/591 C11 staining in CCF cells treated with vehicle (DMSO), TMZ (100 µM) and FGFR1i (PD166866, 5 µM) for 72 hours. (M) Fold change of ferroptosis-related genes upregulated (red) or downregulated (blue) by TMZ treatment *versus* the combination of TMZ plus FGFR1i. (N) Quantification of cell death as determined by Annexin V staining of CCF cells co-treated with TMZ and FGFR1i either in the presence or the absence of the apoptosis inhibitor Q-VD-OPH (QVD, 20 µM) or the ferroptosis inhibitor ferrostatin-1 (Fer-1, 5 µM) for 72h. Graphs show mean values ±SEM (n=3 biologically independent experiments). \**p* < 0.05 (ANOVA *post hoc* Bonferroni test).

### FGFR1i synergizes with TMZ to prevent tumor growth in a preclinical *in vivo* model

To validate the potential capacity of the dual treatment with FGFR1i plus TMZ as a therapeutic approach, we resort to mouse models using xeno-implants. Although we initially essayed an orthotopic model by intracranial injection of GBM cells in the dorsal striatum of nude mice, these cells were not able to successfully implant in the brain (data not shown), and therefore we used subcutaneous xeno-implants in the flank of nude mice. This model is more limited, as we cannot assess potential physiological limitations of the treatment related to the blood–brain barrier. Nevertheless, we were still able to validate the capacity of FGFR1 inhibitors to target tumor cells and to synergize with TMZ in physiological conditions. Thus, nude mice were implanted in the flank with 3 x 10^6^ CCF cells using Matrigel system. Once tumors reached 100 mm^3^, mice were treated with TMZ and/or FGFR1i by intraperitoneal injection, and tumor growth was followed twice a week (Figure 6A). Animal health was monitored by body weight measurement along the study. Body weight showed no changes neither with TMZ nor FGFR1i treatment, alone or in combination (Supplementary Figure 6A). No effects were observed in mice behaviour nor wellness (data not shown), indicating that this therapeutic approach is well tolerated in preclinical setting. To further validate the safety of the dual treatment, we cultured mouse primary astrocytes and treated them *ex vivo* with TMZ and/or FGFR1i. Presence of primary astrocytes in the culture was confirmed through GFAP staining (Supplementary Figure 6B). As shown in Supplementary Figure 6C, FGFR1 inhibition, with or without TMZ, did not induce cell death in primary astrocytes as measured by annexin-V/PI staining. The treatment did not affect neither ERK nor p53 signaling in primary astrocytes, in contrast to GBM cells (Supplementary Figure 6D). These results confirmed that dual treatment with TMZ and FGFR1i is a safe approach against GBM tumors. Importantly, both TMZ or FGFR1i treatment alone caused a partial decrease in tumor growth, but this decrease was more pronounced in mice receiving the dual treatment (Figure 6A). Indeed, FGFR1i had a further synergistic effect towards tumor regression in TMZ-treated mice (Figure 6B and Supplementary Figure 6E), confirming our results using cultured cells. It was somehow surprising to observe that, despite the strong TMZ resistance observed in these cells *in vitro*, TMZ treatment had a significant effect on tumor growth inhibition *in vivo*, yet less pronounced than the dual treatment. This TMZ-mediated effect is likely a cytostatic effect, rather than cytotoxic, as inferred by cell cycle arrest induced by TMZ in these cells. Histological (H&E) analysis confirmed massive regression in tumors treated with FGFR1i plus TMZ, with necrotic areas and reduced cellularity (Figure 6C). To confirm the molecular mechanism of the therapeutic approach in the implanted tumor, we performed immunohistochemical analysis. As shown in Figure 6D-F and Supplementary Figure 6F, we confirmed a marked reduction in FGFR1 phosphorylation in FGFR1i plus TMZ treated cells, which correlated with the decrease in Ki67 staining (Figure 6F). It is worth noting that FGFR1 inhibition slightly restored Ki67 staining in TMZ-treated tumors, suggesting that FGFR1i partially restored cell cycle *in vivo* (as observed *in vitro*). Apoptotic cells were detected almost exclusively in the tumors cotreated with TMZ and FGFR1i (Figure 6D) supporting the cytostatic effect of TMZ treatment alone. These results confirmed the efficacy of the dual treatment to block tumor growth through the inhibition of FGFR1 pathway. Further confirming the molecular mechanism involving p53 induction in TMZ resistance, immunoblot analysis showed an increase in p53 levels in TMZ-treated tumors, which was decreased upon FGFR1i co-treatment (Figure 6G). Of note, and contrary to what we observed in cultured cells, ERK signaling was induced in response to TMZ, as shown both in tumor section (Figure 6D) and immunoblot (Figure 6G). FGFR1i treatment demonstrated a strong capacity to prevent this TMZ-mediated activation of ERK pathway (Figure 6D, 6G) and confirmed the sufficiency of FGFR1 inhibition to block ERK signaling in this genetic background *in vivo*.

**Figure 6.**
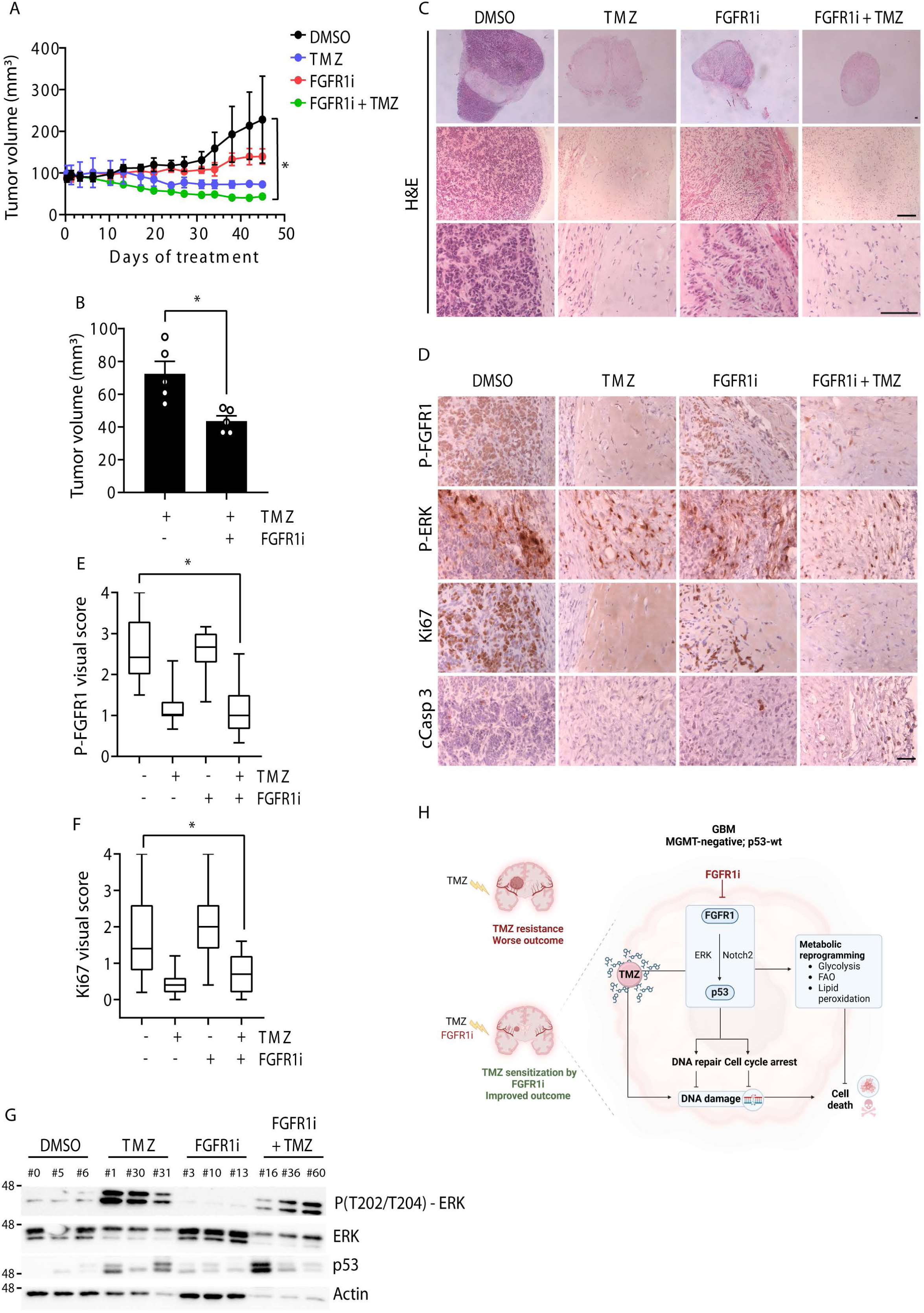
FGFR1 inhibition and TMZ co-treatment efficiently induced tumor regression in a xenograft tumor model. (A) Evolution of xenograft tumor volume during time in mice treated as indicated (n= 5 mice per treatment). Treatments started when tumors reached 100mm^3^. (B) Bar graph representation of tumor volume differences at endpoint between TMZ treatment and the combination of TMZ with FGFR1i. (C) Representative histological images of tumor sections from the xenograft tumors with the different treatments, stained with haematoxylin and eosin (H&E). Scale bars represent 100μm. (D) Representative images of the immunohistochemistry on sections from xenograft tumors at endpoint for the indicated markers and counterstained with haematoxylin. Scale bar represents 50μm. (E, F) Visual score analysis of images as in (D) for P-FGFR1 (E) and Ki67 (F) staining. The upper and lower limits of the boxes represent quartiles, the line within the boxes indicates the median and the whiskers show the extremes (n ≥ 15 images per treatment). (G) Immunoblot for ERK activity (ERK phosphorylation) and p53 from 3 different xenograft tumors treated as indicated. (H) Scheme of the working model representing the signaling and metabolic reprograming associated with TMZ resistance and controlled by FGFR1. Figure 6H was created with BioRender.com.

Altogether, our mouse model confirmed that FGFR1i in combination with TMZ safely targets FGFR1-positive GBM cells *in vivo*, and therefore supports this strategy as a potential approach against this aggressive type of brain tumors to be confirmed in patients.

## DISCUSION

Despite the efforts in basic and clinical research, GBM continues to represent an unmet need in medicine and health care. The current standard of care, including TMZ treatment, remains mostly palliative, with strong treatment resistance problems in almost all patients who do not receive satisfactory solutions from the available therapies [14], [29]. Two working approaches are in progress to provide with new alternatives: i) exploring different strategies to induce GBM cell death and tumor regression; and ii) addressing recurrent resistance to current therapies. The work presented herein is in line with both approaches. We propose FGFR1 receptor activity as a novel therapeutic target in resistance to TMZ. Our results suggested that GBM patients with an active FGFR1 had worse prognosis than GBM patients with low or undetectable FGFR1 activity. FGFR1 was the main regulator of cell signaling and cell growth through MAPK/ERK pathway. Through a multi-omics (transcriptomics, metabolomics and lipidomics) analysis, we identified a deep remodeling in GBM metabolism associated with TMZ resistance. Notably, this metabolic remodeling was strongly controlled by FGFR1, as FGFR1 inhibition prevented this metabolic rewiring, and importantly, restored TMZ sensitivity. Noteworthy, none of the elements involved in the global program induced by TMZ and controlled by FGFR1 were sufficient to explain individually the resistance to therapy, suggesting that FGFR1 synergism with TMZ operates as a consequence of blocking the whole resistance program. This conclusion underscores the importance of targeting FGFR1, but no other of its downstream elements, to overcome TMZ resistance in GBM.

It is important to highlight that our results showed that the efficacy of FGFR1 inhibition greatly depended on the level of activity of this receptor and on the genetic background.

Coherently, only FGFR1-positive GBM cells responded to inhibition. Supporting FGFR1 role in GBM chemoresistance, a recently published work found the non-coding microRNA miR-3116 to target FGFR1 expression and to impact on TMZ sensitivity in GBM cells [30]. In addition to that, this approach only showed efficacy in a MGMT-negative, p53 WT context. MGMT is the enzyme that eliminates DNA damage induced by TMZ, and as such, its activity is a main mechanism in GBM tumors to induce TMZ resistance. However, in those tumors presenting MGMT-promoter hypermethylation and therefore no MGMT expression (MGMT-negative), resistance to TMZ greatly depends on a p53-mediated DNA repair program, including cell cycle arrest (reviewed in [31]). Our results indicated that this protective p53 program was controlled by the FGFR1 pathway. Indeed, inhibition of FGFR1 abolished the response of p53 to TMZ and the subsequent cell cycle arrest. Therefore, both p53-dependent program and metabolic rewiring involved in TMZ resistance required an active FGFR1 signaling pathway, constituting the molecular basis of the synergistic effect observed when combining TMZ treatment with FGFR1 inhibition, described in the present work (Figure 6H). Our results, thus, also highlighted the importance of an appropriate GBM patient stratification. At the moment, only pan-FGFR inhibitors are in clinical trials for specifically recurrent GBM harboring FGFR activating mutations or the FGFR-TACC oncogenic fusions (https://clinicaltrials.gov/). However, this type of FGFR alterations occurs in a very small subset of human GBMs (3%) [32]. More than 60% of GBM patients harbors an unaltered TP53 gene [33], and our results, yet in a limited patient cohort, showed a 22% of GBM samples with high/very high FGFR1 activity. Hence, we postulate that a wider GBM patient population would benefit from specific FGFR1 inhibition provided that a good patient stratification is implemented.

The relationship between FGFR1 and Notch2 signaling in GBM remains elusive after this work. We did not observe a direct biochemical interaction between both proteins (data not shown), as initially hypothesized. Still, FGFR1 inhibition deeply abrogated Notch2 signaling, suggesting that Notch2 operates downstream of FGFR1 in GBM. This result contrasted with what was described in NSC, in which expression levels of Notch2 impacted FGFR1 signaling [20], a result not observed in GBM cells. Still, the role of Notch2 in FGFR1-mediated chemosensitivity was not substantiated in this work. It is likely that FGFR1 and Notch2 coordinate the signaling and metabolic program during TMZ resistance, but certainly FGFR1 exerted a dominant role with respect to other elements, including Notch2.

Finally, our work highlights the role of metabolic reprogramming during therapy resistance. The dual treatment (TMZ + FGFR1i) increased lipid peroxidation associated with ferroptosis. Still, our results indicated that blocking specific metabolic pathways was not sufficient to restore TMZ resistance, once again reflecting that TMZ resistance did not operate through one exclusive mechanism, but through a whole signaling and metabolic rewiring, which is controlled by FGFR1 signaling. All these results strongly point toward the high potential of using FGFR1 inhibitors as co-treatment to prevent TMZ resistance in GBM patients.

## METHODS

### Human tumor samples

A total of 27 paraffin-embedded human GBM samples were provided by the Biobank of the Anatomy Pathology Department (record number B.0000745, ISCIII National Biobank network) of the MD Anderson Cancer Centre Madrid, Spain. These samples were recruited between since 2016 and 2021, with an age of 70.48 ± 14.5, comprising 17 men and 9 women, were diagnosed with grade IV astrocytoma/glioblastoma in accordance ith the pathological criteria, irrespective of treatment or other parameters. All patients underwent surgical and/or chemotherapeutic treatment in accordance with the prevailing standards of care. This study was conducted in accordance with the ethical standards set forth in the Spanish legislation (*Ley de Investigacion Organica Biomedica*, 14 July 2007) and was approved by the ethical committees of the MD Anderson Cancer Centre Madrid, Spain. All patients consented to the study, having been provided with written information and having given their written informed consent.

### Cell culture

T98G (ATCC® CRL-1690™), CCF (ATCC® CRL-1718™), LN18 (ATCC® CRL-2610™), and LN229 (ATCC® CRL-2611™) cells were obtained from the American Type Culture Collection (ATCC); A172 (catalogue number 300108) cells were obtained from Cell Lines Services (CLS); and U87 cells were kindly provided by Prof. Abelardo López (CABIMER, Seville, Spain). All the cell lines were cultured and maintained in DMEM (D6546, Merck) supplemented with 10% of Fetal Bovine Serum (FBS), 2mM of glutamine, penicillin (100IU·ml^-1^) and streptomycin (100µg·ml^-1^) at 37°C, 5% CO_2_ in a humidified atmosphere. All the cell lines were authenticated and regularly tested for mycoplasma contamination. Where indicated, the following compounds were added to the culture medium: 4 µM PD173074 (sc202610, Santa Cruz Biotechnology), 2.5 or 5 µM PD166866 (PZ0114, Merck), 100 µM TMZ (T2577, Merck), 10 µM DAPT (sc-201315, Santa Cruz Biotechnology), 1 µM CB-839 (5337170001, Meck), 4 µM Etomoxir (5337170001, Merck), 20 µM Q-VD-Oph (SML0063, Merck), and 5 µM ferrostatin-1 (SML0583, Merck).

### Mouse primary astrocytes from neurosphere differentiation

The brains of 10 animals were collected from P3 littermate pups. Each brain was cut in half and the two striata dissected and mechanically dissociated in ice-cold DMEM-F12 medium (D6421; Merck). After centrifugation, cell pellet was resuspended in neurosphere medium (NSM) consisting of DMEM-F12 supplemented with 2mM glutamine, B27 (17504044, Thermo Fisher), EGF (20ng/ml) (130-097-749, Miltenyi Biotec), and antibiotics penicillin (100IU/ml) and streptomycin (100 µg/ml). Cells were plated in a T75 flask, and cultured at 37°C in humidified 5% CO_2_ with half of the medium refreshed every 3 days. After approximately 10 days, neurospheres were treated with accutase (A6964, Merck) for 10 min at RT, and dissociated cells were plated in a T75 flask with NSM to obtain second-generation neurospheres with medium refreshed every 3 days. After reaching the appropriate size, secondary neurospheres were induced for astrocyte differentiation. After neurosphere dissociation with accutase, cells were seeded in T75 flask with DMEM-low glucose (D5546, Merck) supplemented with 10% FBS. After 2-week differentiation, astrocytes were collected with accutase and seeded for the experimental conditions.

### Immunofluorescence

Astrocyte cultures were analysed by immunofluorescence for the expression of the astrocytic marker GFAP (Glial fibrillary acidic protein). Cells were cultured at 6·10^4^ cells/ml on poly-L-lysine coated coverslips in 24-well plates. Cells were fixed with paraformaldehyde 4% (sc-281692, Santa Cruz Biotechnology) for 10 min, permeabilized with 0.05% Triton-X100 (T9284, Merck) for 10 mins, and incubated with 5% BSA for 30 min to block non-specific sites (04-100-812-D; Euromedex). Incubation with anti-GFAP antibody (80788, Cell signalling) 1:200 in blocking solution for 2h at RT, was followed by 3x rinse with PBS, incubation with donkey anti-rabbit secondary antibody at 1:200 (Cy3, 711-165-152, Jackson Immunoresearch) for 1h in the dark and mounted with ProLong Gold Antifade with DAPI (P36941, Molecular Probes). Images for astrocyte quantification were taken with a fluorescence microscope Zeiss Apotome 3.

### DNA damage measurement

Cells were seeded in triplicate at 3·10^4^ per well in 96-well plates (655096, Greiner Bio-One) with the respective treatments for 72h. Thereafter, cells were fixed with 4% paraformaldehyde for 10 mins at RT (sc-281692, Santa Cruz Biotechnology) and permeabilized with 0.5% Triton-X100 for 10 mins. Non-specific binding was blocked by incubating with 3% BSA for 30 min at RT. Finally, cells were incubated with the primary antibody γH2AX 1:2000 (05-636, Merck), overnight at 4°C. After three washes with PBS, cells were incubated with Alexa Fluor™ Goat Anti-mouse IgG 488 (A28175, Invitrogen), for 1 hour at RT in the dark. Finally, cells were stained with DAPI for nuclear staining. Samples were automatically imaged using the High-Content Microscopy ImageXpress (Molecular Devices), and analysed with MetaXpress software.

### Immunohistochemistry

Paraffin-embedded samples were submitted to standard immunohistochemistry (IHC) protocol with rabbit anti-phospho-FGFR1 (Tyr653/Tyr654) polyclonal antibody, 1:50 (44-1140G, Thermo Scientific), mouse anti-Ki67, 1:100 (550609, BD Pharmingen), rabbit anti-cleaved Caspase 3, 1:100 (9664, Cell Signalling), and rabbit anti-phospho-ERK, 1:100 (4370; Cell Signalling). Anti- rabbit and anti-mouse biotinylated secondary antibodies were used for 1h at RT (21537 and 21538 respectively, Merck), followed by vectastain ABC reagent (PK4000, Vector Laboratories). DAB reaction kit (SK-4100, Vector Laboratories) was used to visualise the peroxidase reaction product. Samples without the primary antibody were used as a negative control. All the slides were counterstained with haematoxylin 1:3 diluted. For haematoxylin and eosin staining (H&E) paraffin-embedded samples were submitted to standard H&E staining protocol. Images were obtained with a Leica microscope and Leica digital camera. Visual score analysis was performed from 5 independent blinded analyses of ≥ 12 images/sample. Immunostaining was classified using a score 0 to 4, calculated by multiplying staining intensity (0 for no staining, 1 for light/weak, 2 for intermediate intensity, 3 for strong, and 4 for very strong). Tissue spots that were missing, damaged or contained staining artefacts were excluded from the analysis.

### Western blot analysis

Cells from a 6 cm plate or a 6-well plate were rinsed with PBS and proteins were extracted using RIPA lysis buffer (50 mM Tris, 150 mM NaCl, 1 mM EGTA, 1 mM EDTA, 0.1% (v/v) SDS, 1% (v/v) Triton X-100, 0.5% (w/v) sodium deoxycholate, pH 7.4), containing phosphatase inhibitors (P0044 and P5726, Merck) and a cocktail of protease inhibitors (P8340, Merck). Protein concentration was quantified by the Bradford assay (BCA kit, 23225, Thermo Fisher). Lysates were resolved by SDS-PAGE electrophoresis and transferred to nitrocellulose membrane (0.2μm, 1620112, BioRad) using the Trans-Blot Turbo system (Biorad). Blocking for 1h in 5% BSA (in TBS with 0.01%Tween-20), followed by overnight incubation with the primary antibody (Supplementary Table 1) at 4°C. Horse-radish peroxidase conjugated secondary antibodies (Supplementary Table) were used for 1h at RT, and bands were detected using ECL reagent (1705061, Biorad). Images acquired using the ChemiDoc-MP imaging system (Biorad).

### Flow cytometry

For cell death detection, cells were harvested by trypsinization and resuspended in DMEM. Cells were stained with annexin V and propidium iodide (PI) from the Annexin V-FITC Early Apoptosis Detection Kit (ANXVKF, Immunostep) following the manufacturer’s protocol. For cell cycle analysis, cells were harvested by trypsinization and fixed in cold 70% ethanol added dropwise while vortexing for 30 min at 4°C. Cells were then rehydrated with PBS for 2 min and rinsed twice with PBS. Cells were stained with a solution containing 1μg/ml PI (P4170, Merck) and 100μg/ml RNAse (10109134001, Merck) for 15 min at 37°C. Data was acquired on a BD-Bioscience LSRFortessa X-20 flow cytometer and analyzed with BD Diva software.

### Real-time PCR

Total RNA was isolated using TRIzol reagent (12044977, Thermo Fisher) and quantified with the NanoDrop™ 2000 (Thermo Fisher Scientific). A total of 1 μg of RNA was retrotranscribed using GoScript Reverse Transcription System (A5003, Promega). Real-time quantitative PCR (qPCR) was performed in duplicate using Step-One Plus Real-Time PCR System (Applied Biosystems), by mixing 1ng cDNA with specific primers and SYBR Green Supermix (1725121, BioRad). Specific primer sequences (Merck) are listed in the Supplementary Table 2.

### Small interfering RNA transfection

Cells were seeded at 4·10^5^ cells per plate in a 6 cm plate. After 24h, siRNA transfection was performed using 10 nM of ON-TARGETplus^TM^ siRNA (Dharmacon^TM^, Horizon) and INTERFERin^R^ siRNA transfection reagent (PPLU101000016, POLYplus transfection) following the manufacturers’ instructions. ON-TARGETplus Non-targeting Control siRNAs were also used. After 48h of transfection, cells were treated for the different conditions and proteins or total RNA were extracted.

### RNA-seq analysis

Total RNA from CCF cells was extracted using the E.Z.N.A. HP Total RNA frit (R6812-01, Omega Bio-Tek). Libraries were prepared using the Illumina Stranded mRNA Prep ligation, and the IDT for Illumina RNA UD Index SetA ligation (20040532 and 20040553, respectively; Illumina) following manufacturer’s instructions. Sequencing was performed employing the Flow Cell NovaSeq 6000 SP reagent kit v1.5 (200 cycles) 2x100bp at the Genomic Unit of CABIMER (Seville, Spain). Three independent biological replicates of each condition were sequenced. RNA-seq samples were aligned to the hg38 human reference genome (*Homo sapiens*). Genome index was generated with *buildindex* function and alignment with the *align* function, both from Bioconductor Rsubread package [34], using type = 0, TH1 = 2 y unique = TRUE parameters. The.dam files were ordered and indexed using the Rsamtools (v 2.16.0) *sortBam* and *indexBax* functions (Morgan et al, 2020). To assign reads to the UCSC hg38 Known genes, it was used the FeatureCounts function from the Bioconductor Rsubread package, using the options GTF.featureType = “exon”, GTF.attrType = “gene_id” y strandSpecific = 2. The downstream analysis was followed by the differential gene expression analysis, performed using a standard protocol with functions from Bioconductor voon/lima (v.3.34.9) and edgeR (v3.42.4), and the DESeq2 (v1.40.2) packages [35]. Differentially expressed genes with a threshold of non-adjusted *p* value < 0.05 and |Log_2_FC| > 0.5 were selected for further analysis. Heatmaps were performed with *pheatmap* (v1.0.12). Gene Set Enrichment Analysis (GSEA) was analysed using the GSEA v2.0.14 software (GSEA, Broad Institute, Cambridge, MA, USA) for Linux (v4.3.2) with 1000 permutations [35]. Biological processes from functional annotations categories were analyzed with DAVID [36] or with ShinyGO 0.81 [37]. Venn diagrams were performed using Venny 2.1 (https://bioinfogp.cnb.csic.es/tools/venny/index.html).

### Oxygen consumption and extracellular acidification rate measurements

Seahorse technology, using an XF-24 extracellular flux analyser (Agilent), that provides real-time measurements of oxygen consumption rate (OCR) and extracellular acidification rate (ECAR) was applied to determine mitochondrial respiration capacity and glycolysis rate respectively. Cells were seeded at 3.8·10^5^ per plate in 6 well plate and treated for 72h. The day before the experiment, cells were collected by trypsinization and 4·10^4^ cells of each condition were seeded in XFe24 cell culture microplate (Agilent) in DMEM maintaining the treatment, and incubated at 37°C, 5% CO_2_ in humidified atmosphere. After 24h, cells were rinsed and incubated with phenol-red-free DMEM a maximum of 1h at 37°C with atmospheric CO_2_ in a non-humidified incubator in order to eliminate residues of carbonic acid from medium. The cartridge was rehydrated following manufacturers’ instructions. For OCR analysis, changes in concentrations of dissolved oxygen were collected before and after the addition of different inhibitors to the cartridge following manufactureŕs instructions: 1 μM oligomycin (75351, Merck) (port A), 2.5 μM FCCP (Carbonyl cyanide p-(trifluoromethoxy) phenylhydrazone, C2920, Merck) (port B), 1 μM rotenone (R8875, Merck) and 1 μM antimycin A (A8674, Merck) (port C). Three measurement cycles consisted of 2-min mix, 2- min wait, and 4-min measure were carried out. For the glycolytic rate analysis, changes in extracellular free protons were collected before and after the inhibitors 0.5 μM rotenone and 0.5 μM antimycin A (port A), and 50 mM 2-Deoxy-D-glucose (2-DG) (D8375, Merck) (port B). Three measurement cycles consisted of 3-min mix, 2- min wait, and 3-min measure were carried out at basal condition and after each injection. For standardisation purposes, at the end of the experiments the medium was removed, each well was carefully washed twice with 50 μL of PBS and proteins were extracted with 50 μL of RIPA-Hard lysis buffer at RT, and quantified by BSA assay.

### Bodipy assay

For lipid droplet staining, cells were seeded and treated as usual. After 72h of treatment, cells were stained with 2 μM Bodipy FL C12 (D3822, Thermo Fisher) for 30 min at 37°C in the dark. Cells were then harvested by trypsinization and Bodipy staining was assessed by flow cytometry. For Bodipy lipid peroxidation assay, cells were treated as usual. Bodipy C11 581/591 lipid peroxidation sensor (D3861, Thermo Fisher) was added at 4 μM as final concentration and incubated for 30 min at 37°C. Cells were then harvested and analysed by flow cytometry. For both assays, data was analysed with FlowJo v10 (BD Biosciences) and represented as the mean fluorescence intensity (MFI).

### Stable isotope tracing and analysis

For the tracing experiments, cells were treated with TMZ and FGFR1i for 72h, washed twice with PBS and cultured with medium containing the labelled metabolite for 4h. For glucose tracing, DMEM without glucose was supplemented with dialysed FBS (dFBS), 2 mM Glutamine and 25mM 1,2 ^13^C_2_ D-Glucose (CAS 138079-87-5, Cambridge Isotope Lab). For glutamine tracing, DMEM-high glucose without glutamine was supplemented with 10% dFBS, and 2 mM ^13^C- glutamine (CAS 184161-19-1, Cambridge Isotope Lab). For palmitate tracing, DMEM-high glucose without glutamine was supplemented with 10% dFBS, 2 mM glutamine and ^13^C-sodium palmitate (CLM-6059-PK, Cambridge Isotope Lab). On the day of metabolite extraction, cells of each experimental condition were counted to calculate the volume of extraction solvent (1·10^6^ cells/ml solvent). Cells were washed twice with cold PBS and after adding the extraction buffer (50:30:20, v/v/v acetonitrile/methanol/water), cells were scrapped, transferred to a prechilled cold eppendorf and incubated 20 min at -20°C. After vortexing, samples were centrifugated at 21,000 g for 10 min at 4°C, and supernatants transferred to liquid chromatography-mass spectrometry glass vials. Samples were stored at -80°C until further analysis by liquid chromatography - mass spectrometry (LC-MS) as previously described [38]. Briefly, metabolic profiling of the polar metabolome is achieved by using HILIC chromatography coupled with high- resolution mass spectrometry. Separation was performed using an Ultimate 3000 high- performance liquid chromatography (HPLC) system (Thermo Fisher Scientific) on a ZIC-pHILIC guard and analytical column (SeQuant; 150 mm by 2.1 mm, 5 μm; Merck) maintained at 45°C. The gradient started with 20% ammonium carbonate [20 mM (pH 9.2)] (A) and 80% acetonitrile (B), and increased linearly to 80% phase A at a constant flow rate of 200 μl/min. For MS, a Q Exactive orbitrap mass spectrometer (Thermo Fisher Scientific) equipped with electrospray ionization was used. Data was acquired in polarity switching mode with a resolution (RES) of 70,000 at 200 mass/charge ratio (m/z), over a mass range of 75 to 1000 m/z. Minimum automatic gain control (AGC) and maximum injection time were set to 1e6 and 250 ms, respectively. Targeted metabolomics analysis was performed using Skyline software (v21.2.0.369). Extracted ion chromatograms were generated for each compound and its isotopologues using the m/z of the singly charged ions (XIC, +/- 5 ppm) and explicit retention times with 2-min retention time window from our in-house metabolite library, which was generated using reference standards on the same LC-MS method. The ^13^C labelling analysis was determined by quantifying peak areas of all isotopologues of each metabolite by knowing its accurate mass. Labelled isotopologues are defined and represented as M + N, where “M” refers to the metabolite mass, and “N” to the number of ^13^C incorporated.

### Lipidomics

Free fatty acids (FFA) were extracted from biological samples using a mixture of butanol and methanol (Bu: Me) in a ratio of 50:50 containing heptadecanoic acid (C17:0) at a concentration of 1 µM as an internal standard (ISTD). 200µl of the extracts were then evaporated under N_2_ and reconstituted in 30 µl of Bu: Me for subsequent analysis using LC-MS. Analysis of FFA was performed using an Ultimate 3000 HPLC (Thermo Fisher Scientific) coupled to a Q-Exactive Orbitrap mass spectrometer (Thermo Fisher Scientific). 5 µL of sample were injected into the LC-MS instrument and separation was achieved on an Acquity UPLC CSH C18 column (100 × 2.1 mm; 1.7 µm; Waters Corporation) maintained at 55°C. The mobile phases consisted of (A) acetonitrile: water (60:40) containing 10 mM ammonium formate and 0.1% formic acid; and (B) isopropanol: acetonitrile (90:10) containing 10mM ammonium formate and 0.1% formic acid. Gradient elution was used starting at 10% B for 2 min, then increased to 90% at 6 min, and maintained for 1 min, then it was returned to 10% at 8 min and held constant till the end of the run at 10 min. Autosampler temperature was maintained at 4°C. Data was acquired using negative polarity, over the following mass range 150–500 m/z at RES 70,000. Peak identity was confirmed by comparison to reference standards and mathematical calculations using regression curves obtained by plotting the retention times of fatty acids of known chain lengths and number of double bonds. Overall, the method measured 34 FA with chain lengths between 10-26 and double bonds between 0-6. Data analysis was performed using Skyline software (v 21.2.0.369). A transition list containing the accurate m/z and the explicit RT (±2-min retention time) window for each FFA was generated using our in-house FFA library. All peaks were integrated, normalised to total cell number and peak area ratios were calculated relative to the ISTD. Data was plotted for peak area ratios of each FA.

### Xenograft mouse model

Animals were housed in pathogen-free conditions at the Animal Facility in CABIMER. The project received the agreement from the Ethics Committee on Animal Experimentation of *Junta de Andalucia* (Spain) (Agreement number: 28/11/2023/105). 7-week-old female NOD.Cg-Prkdc^scid^ ll2rg^tm1Wjl^/ SzJ immunodeficient mice were randomly assigned to four different groups (5 mice per group). A total of 3·10^6^ CCF cells in 150 μL mixture of medium and 66% Matrigel (corning TM Matrigel GFR Membrane Matrix, 354230, Corning) were subcutaneously injected into the right dorsal flank of the mouse. Tumor growth was assessed with a calliper twice weekly. Once the average tumor volume reached 100 mm^3^, intraperitoneal drug treatment was initiated. PD173074 was administered at 20 mg/ kg/ day, and TMZ at 40mg/ kg/ day, three times per week using a volume of 150 μL.

### Bioinformatic analysis

Box plots for gene expression levels, Kaplan-Meier overall survival curves and survival heatmap in human GBM tumors vs normal tissue samples were calculated from the TCGA (Pan Cancer Atlas) and the GTEx databases using the Gene Expression Profiling Interactive Analysis (GEPIA2) online tool [39]. Survival heatmap estimated using Mantel–Cox test. Data analysis of FGFR1 and Notch2 mRNA expression in TMZ treated and non-treated patient cohorts was performed using GBM patient data from the TCGA database obtained through the cBioportal website, clinical and genomic data from the dataset gbm_tcga_pan_can_atlas_2018. GraphPad Prism software 9 (GraphPad Software, San Diego, CA, USA) was used for analysis, statistics and representation. Two-way ANOVA followed by Bonferroni’s comparison were used to evaluate the statistical difference of the results. Statistical significance was estimated when *p<0.05*. Clinical data analysis based on the phospho-FGFR1 immunohistochemistry of the human GBM samples was performed with R and Bioconductor software. Kaplan-Meier curve was created with the ggplot2 package. Relapse-free survival was defined as the time from surgery until the first instance of recurrence. To run this analysis samples were grouped into FGFR1-negative or FGFR1-positive according to FGFR1 phosphorylation levels.

### Statistics

The results are expressed as mean values ±SEM of at least 3 biologically independent experiments, otherwise indicated. One-way ANOVA followed by Bonferroni’s comparison as a *post hoc* test was used to evaluate the statistical difference of the results. Statistical significance was considered when *p*<0.05. Immunoblots and flow cytometry fluorescence intensity plots are representative of at least three independent experiments.

## ACKNOWLEDGEMENTS

The authors thank the support from the technical services at CABIMER, and the Chromatography Unit of CIC-Cartuja. Research in RVD lab and MT were supported by Grant PID2021-124251OB- I00 and Grant RED2022-134927-T funded by MICIU/AEI/10.13039/501100011033 and by ERDF/UE. LZ was recipient of a predoctoral FPU contract (FPU19/04914) from the Spanish Ministry of Universities. IGLC was recipient of a predoctoral contract (PREDOC_00345) funded by PAIDI, Junta de Andalucia. ARB was recipient of a predoctoral FPI contract (PRE2022-101357) funded by MICIU/AEI /10.13039/501100011033 and FSE+. JMSP and VCG were supported by the Institute of Health Carlos III, co-funded by *Fondos FEDER* (CD23/00104 and CP19/00046). Figure 6H and Supplementary Figure 4H were created with BioRender.com.

## AUTHOR CONTRIBUTIONS

MT and RVD conceived the project. LZ, GVH, MT and RVD designed experiments. LZ, IGLC, MMH, ARB, KMR, JMSP, MCC, VCG, PSM and MT performed experiments. LZ, ESE, SSO, GMB and JCR provided computational resources and performed computational analysis. LZ, IGLC, ESE, SSO, GMB, JCR, GVH, MT and RVD analysed data. GMB, JCR, GVH and RVD secured funding. LZ, MT and RVD wrote the manuscript. All authors read and approved the final version of the manuscript.

## COMPETING INTERESTS

The authors declare no competing interests.

---

**Supplementary Figure 1.**
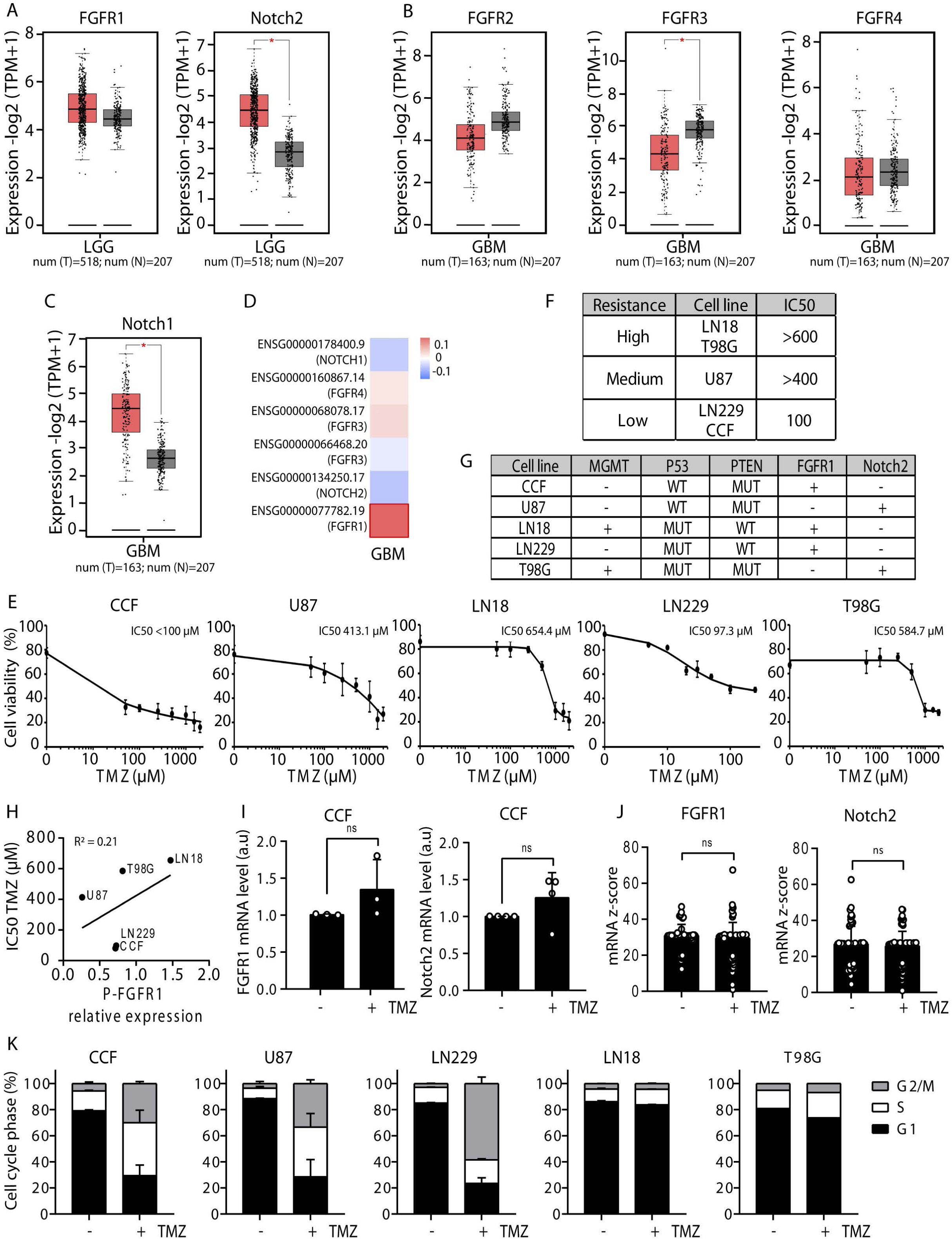
FGFR1 activation correlates with poor prognosis in GBM patients. (A) Box plot for FGFR1 and Notch2 gene expression in low grade glioma (LGG) tumors (T, red) and normal tissue (N, grey) based on tumor and normal samples from TCGA and GTEx databases. (B, C) Box plot for FGFR2, FGFR3, and FGFR4 (B) and Notch1 (C) gene expression in GBM (T, red) and normal tissue (N, grey) analysed as in (A). (D) Survival heatmap for GBM cohort from TCGA and the GTEx datasets showing the survival contribution of the indicated genes. (E) Cell viability curves to establish TMZ IC50 for the indicated GBM cell lines. (F) Classification of GBM cell lines for the resistance to TMZ treatment according to the IC50 analysis in (E). (G) Classification of GBM cell lines according to their genetic status and the activation of FGFR1 and Notch2 receptors. MGMT unmethylated (+) and methylated (-); wildtype (WT) or mutated (MUT): low (-) and high (+) activation. (H) Scatter plot and correlation coefficient between phospho-FGFR1 (P-FGFR1) expression and TMZ IC50 in GBM cell lines. (I) mRNA levels for FGFR1 and Notch2 upon TMZ treatment in CCF cells, assessed by qPCR. (J) mRNA expression analysis for FGFR1 and Notch2 from RNA-seq clinical data from the TCGA Pan-Cancer Atlas GBM dataset. (K) Stacked bar graphs showing the percentage of cells at the different stages of cell cycle upon TMZ treatment of GBM cell lines.

**Supplementary Figure 2.**
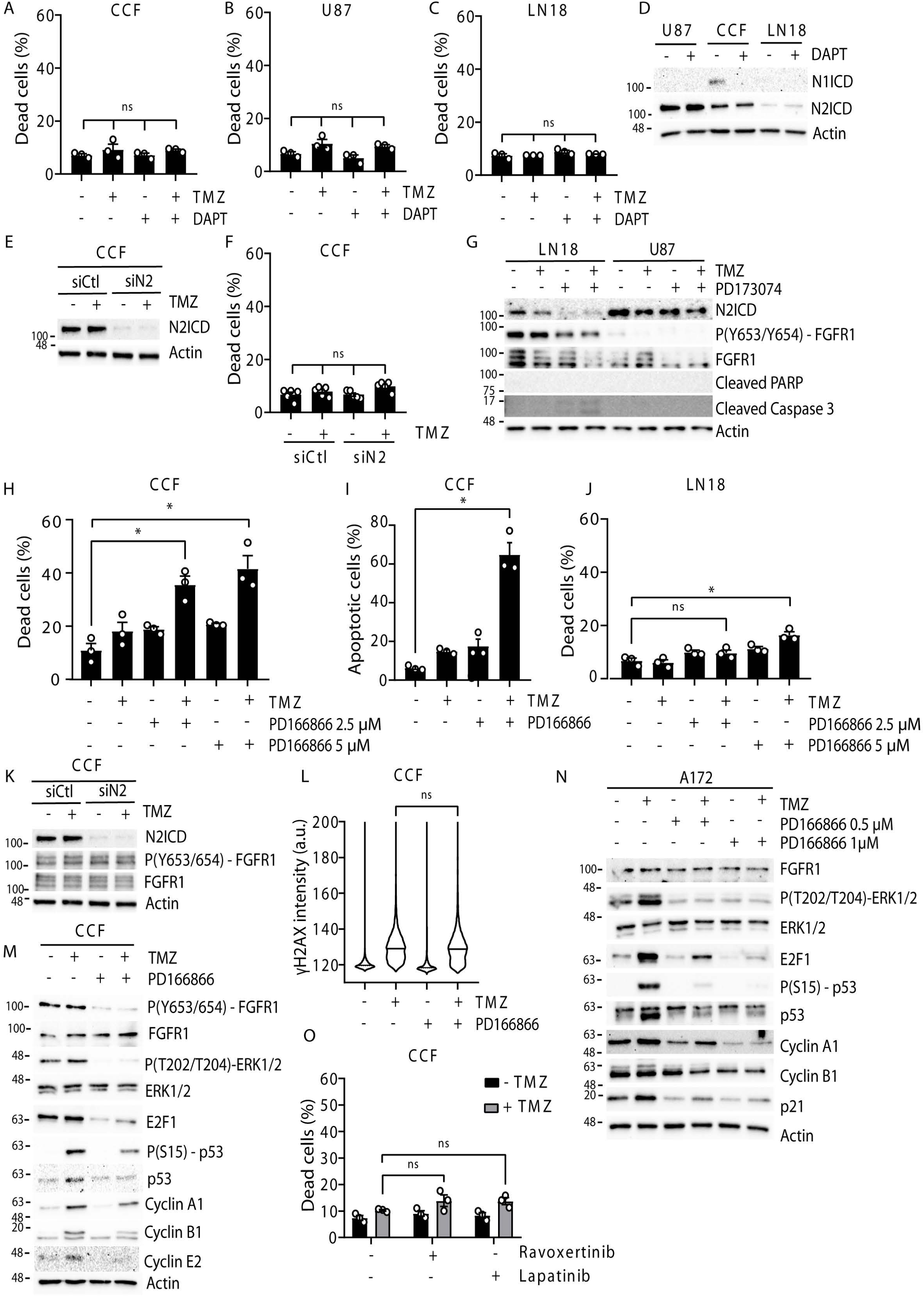
FGFR1 inhibition restores TMZ sensitivity. (A, B, C) Percentage of dead cells estimated by trypan blue viability assay for CCF, LN18 and U87 treated with vehicle (DMSO), TMZ (100 µM) and DAPT (10µM) for 72 hours. (D) Immunoblot analysis of Notch signaling activation (N1ICD and N2ICD) in U87, CCF, LN18 treated with DAPT (10µM) for 72 hours. (E) Immunoblot for Notch2 activation levels (N2ICD) after Notch2 silencing using siRNA and treated with TMZ (100µM) for 72 hours in CCF. (F) Percentage of dead cells by trypan blue assay for CCF cells after Notch2 silencing and TMZ treatment as in (E). (G) Immunoblot of apoptotic markers (cleaved Caspase3 and cleaved PARP) in LN18 and U87 cells incubated with TMZ (100 µM) and the FGFR1i (PD173074, 4 µM) for 72 hours. (H, I) Percentage of dead cells estimated by trypan blue viability assay (H) and flow cytometry with annexin V / PI staining (I) in CCF cells treated with TMZ (100 µM) and FGFR1i (PD166866 5µM) for 72 hours. (J) Percentage of dead cells estimated by trypan blue in LN18 and U87 treated as indicated for 72 hours. (K) Immunoblot for FGFR1 expression and activity (FGFR1 phosphorylation) upon CCF treatment as described in (E). (L) DNA damage quantification of CCF cells by γH2AX expression using MetaXpress analysis upon treatment with TMZ (100 µM) and FGFR1i (PD166866 5µM) for 72 hours. (M) Immunoblot for FGFR1 activation (FGFR1 and ERK phosphorylation), and cell cycle markers (E2F1, p53 and cyclins) in CCF cells treated with TMZ (100 µM) and FGFR1i (PD166866 5µM) for 72 hours. (N) Immunoblot for FGFR1 activation and cell cycle markers in A172 treated with TMZ (100 µM) and FGFR1i (PD166866, at the indicated concentration) for 72 hours. (O) Percentage of dead cells estimated by trypan blue assay in CCF cells treated with ERK and EGFR inhibitors (Ravoxertinib and Lapatinib respectively) combined or not with TMZ for 72 hours. Graphs show mean values ±SEM (n=3 biologically independent experiments). \**p* < 0.05 (ANOVA *post hoc* Bonferroni test).

**Supplementary Figure 3.**
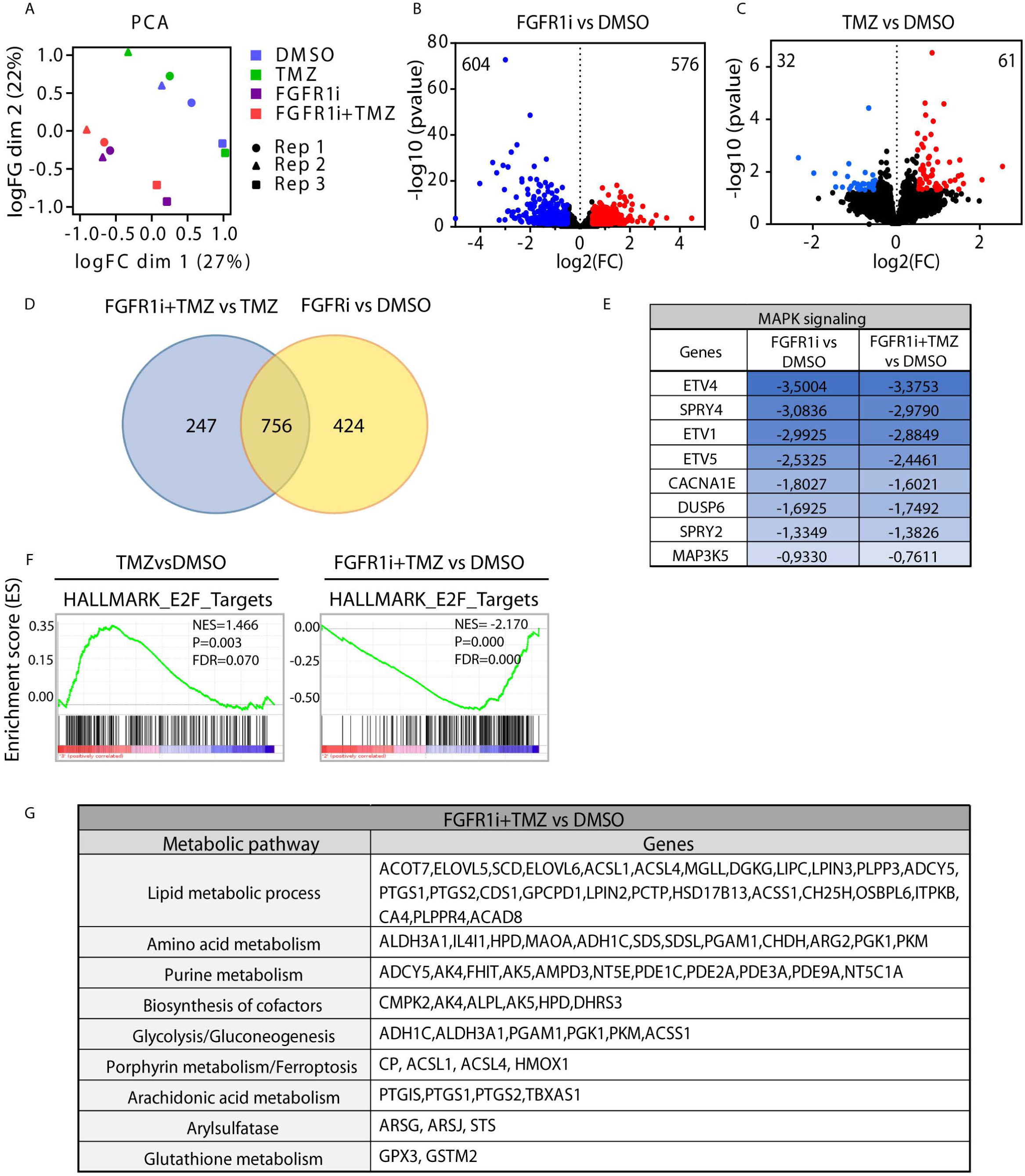
Transcriptomic analysis revealed a metabolic reprograming associated with TMZ resistance. (A) Principal component analysis (PCA) plot showing clustering of the different treatments and replicates from the RNA-seq dataset. (B, C) Volcano plot of the upregulated (red) and downregulated (blue) genes for FGFR1i (B) or TMZ (C) individual treatments. (D) Venn diagram showing the overlapping genes affected upon the indicated treatments as estimated from RNA-seq analysis. (E) Genes from the MAPK signaling pathway dysregulated by FGFR1 inhibition as estimated from RNA-seq analysis. (F) GESEA plot for E2F target genes in the indicated conditions as estimated from RNA-seq analysis. (G) Metabolic genes dysregulated in the dual treatment TMZ plus FGFR1i organised by the metabolic pathway as shown in Figure 3H.

**Supplementary Figure 4.**
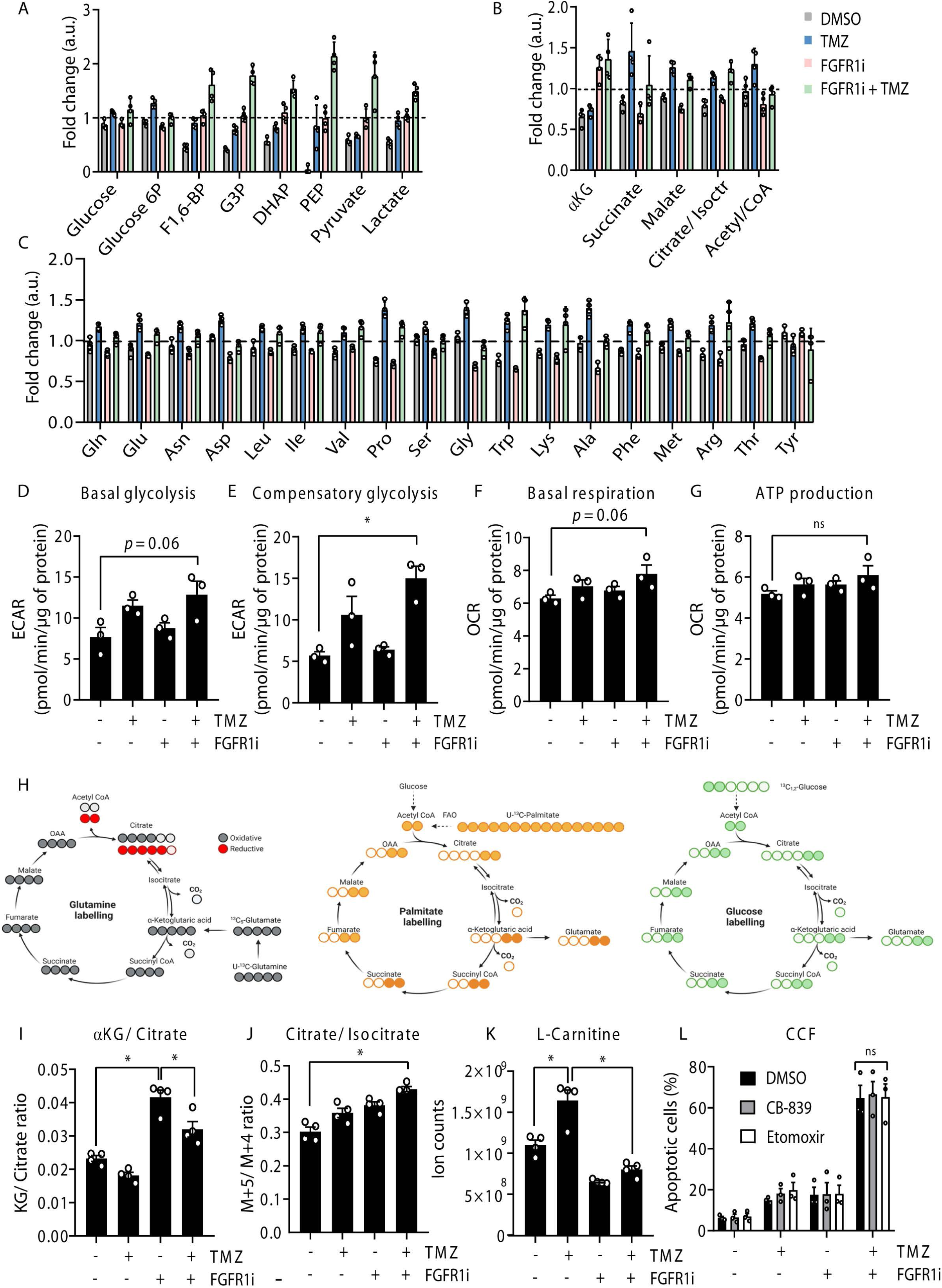
Metabolomic analysis confirmed metabolic rewiring associated with TMZ resistance under the control of FGFR1. (A, B, C) Fold change of glycolytic (A), TCA (B) and amino acid (C) metabolite levels assessed by LC-MS analysis in CCF cells treated with vehicle (DMSO), TMZ (100µM) and FGFR1i (PD166866, 5µM) as indicated for 72 hours. (D, E) Basal and compensatory glycolysis quantification from ECAR analysis in the indicated conditions. (F) Basal respiration quantification from OCR analysis in the indicated conditions. (G) ATP production quantification from OCR analysis in the indicated conditions. (H) Schematic overview to illustrate the fate of the stable isotope-labelled U-^13^C-Glutamine (grey/red), U-^13^C-Palmitate (yellow) and U-^13^C_1,2_-Glucose (green) in the TCA. Carbon atoms are represented by circles. (I) αKG/citrate ratio from CCF cells treated as indicated and labelled with U-^13^C-Glutamine. (J) M+5-citrate/ M+4-isocitrate ratio quantification under the same conditions as in (I). (K) L-carnitine levels in CCF cells treated as indicated as estimated from the LC-MS analysis. (L) Apoptotic cell population obtained by flow cytometry of annexin V/ PI staining of CCF cells treated with vehicle (DMSO), TMZ (100 µM) and FGFR1i (PD173074, 4 µM) for 72 hours, either in the presence or the absence of the glutaminolysis CB-839 (1 µM) or FAO inhibitor etomoxir (4 µM). Graphs show mean values ±SEM (n=3 biologically independent experiments). \**p* < 0.05 (ANOVA *post hoc* Bonferroni test). Supplementary Figure 4H was created with BioRender.com.

**Supplementary Figure 5.**
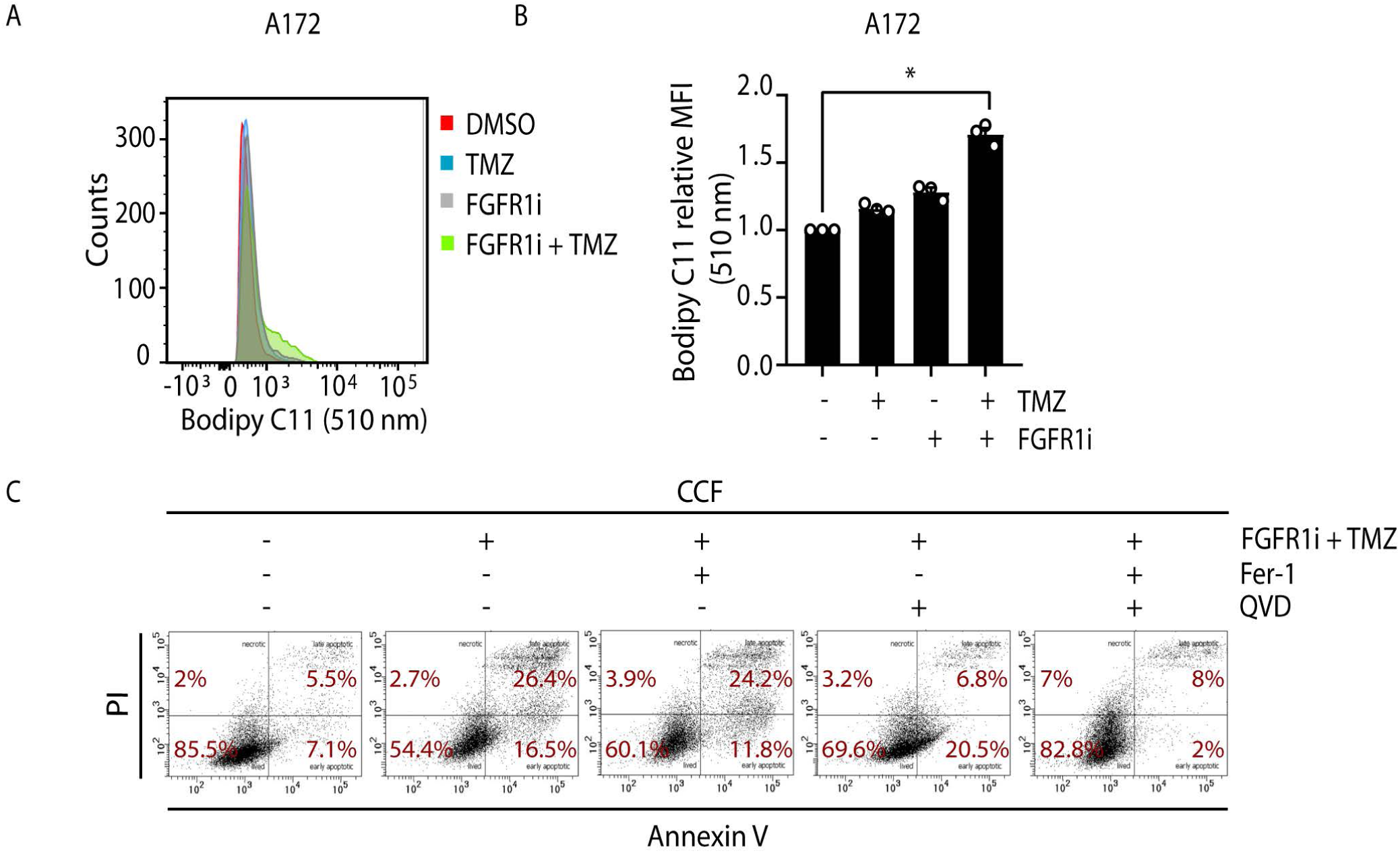
Lipidomic analysis identified lipid profile changes associated with TMZ sensitivity. (A, B) Representative flow cytometry histogram (A) and mean fluorescence intensity quantification (B) of Bodipy 581/591 C11, measured at 510 nm, in A172 cells treated with vehicle (DMSO), TMZ (100 µM) and FGFR1i (PD166866, 5 µM) for 72 hours. (C) Representative flow cytometry dot plots of cell death, measured by Annexin-V and propidium iodide (PI), of CCF cells co-treated with TMZ and FGFR1i either in the presence or the absence of the apoptosis inhibitor Q-VD-OPH (20 µM) or the ferroptosis inhibitor ferrostatin-1 (5 µM) for 72h. Graphs show mean values ±SEM (n=3 biologically independent experiments). \**p* < 0.05 (ANOVA *post hoc* Bonferroni test).

**Supplementary Figure 6.**
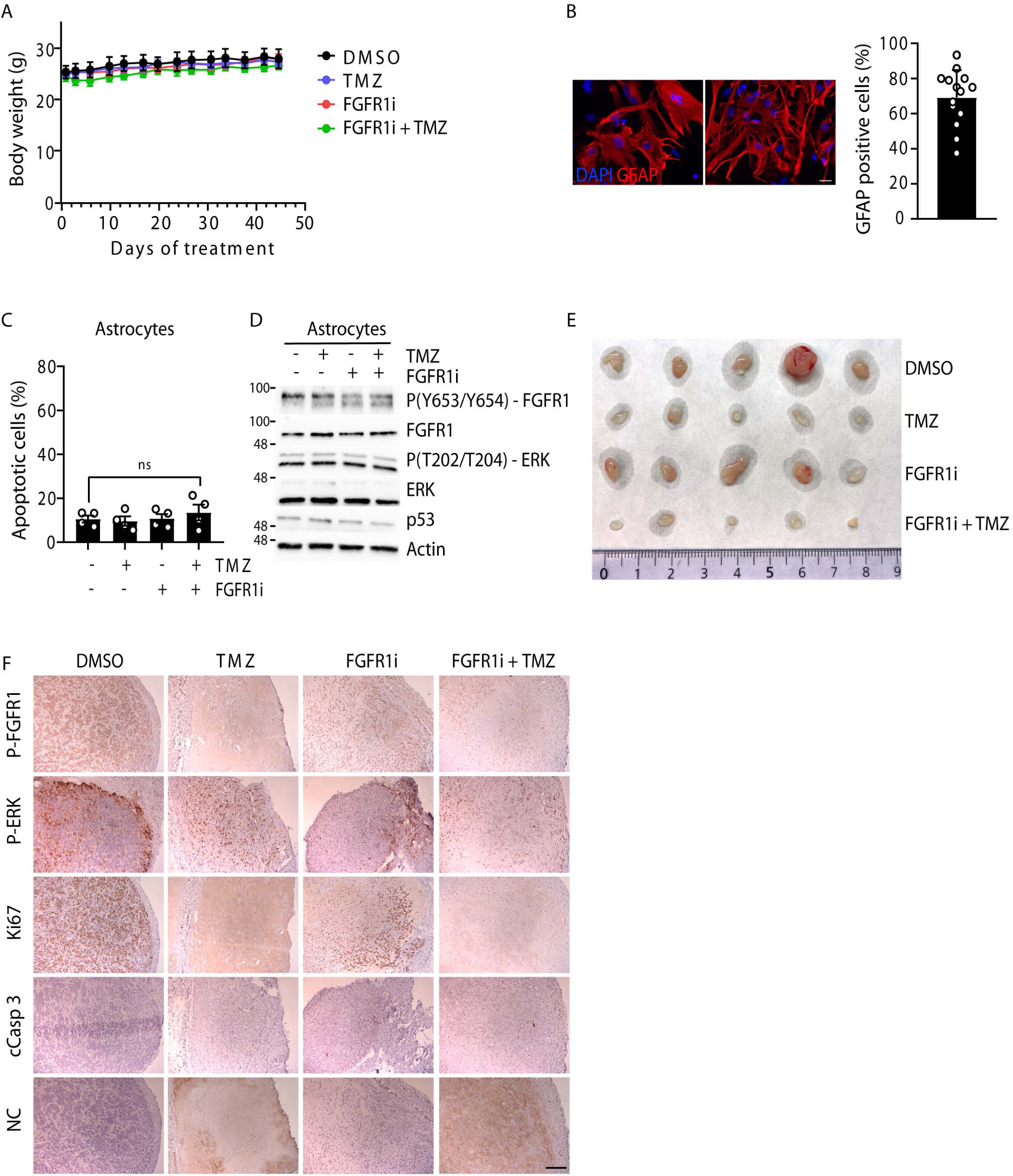
FGFR1 inhibition and TMZ co-treatment efficiently induced tumor regression in a xenograft tumor model. (A) Longitudinal development of body weight of control animals (DMSO) and animals with the different treatments along the experimental period. (B) Two representative immunofluorescence images from primary astrocyte cultures stained with the astrocytic marker GFAP (red), and quantification of GFAP positive cells (n=14 images). (C) Quantification of apoptotic cells by flow cytometry analysis of annexin V/ PI staining of primary astrocyte cultures treated with vehicle (DMSO), TMZ (100 µM) and FGFR1i (PD166866, 5µM) for 72 hours. (D) Immunoblot of FGFR1 (total and phosphorylated), ERK (total and phosphorylated) and p53 in primary astrocyte cultures treated as indicated. (E) Photograph of 5 xenografts tumors observed at endpoint for each condition of treatment. (F) Lower magnification images of histological sections xenografts tumors from each condition of treatment. Scale bar represents 100 µm.

**Supplementary Table 1.** List of antibodies used in this study, indicating target protein, provider, reference, usage dilution and source.

| Antibodies | Provider | Reference | Dilution | Source |
| --- | --- | --- | --- | --- |
| N2ICD | DSHB | C651.6DbHN | 1:1000 | Rat |
| FGFR1 | Sigma-Aldrich | #05-149 | 1:1000 | Mouse |
| P(Y653/Y654) – FGFR1 | Sigma-Aldrich | #06-1433 | 1:1000 | Rabbit |
| P53 | Cell Signaling | #2524 | 1:1000 | Mouse |
| P(S15) – P53 | Cell Signaling | #9284 | 1:1000 | Rabbit |
| P21 | Cell Signaling | #2947 | 1:1000 | Rabbit |
| E2F1 | Cell Signaling | #3742 | 1:1000 | Rabbit |
| Cleaved PARP | Cell Signaling | #5625 | 1:1000 | Rabbit |
| Cleaved caspase3 | Cell Signaling | #9664 | 1:500 | Rabbit |
| β-Actin | Cell Signaling | #4970 | 1:1000 | Rabbit |
| P(T202/T204)-ERK1/2 | Cell Signaling | #4370 | 1:1000 | Rabbit |
| ERK1/2 | Cell Signaling | #9102 | 1:1000 | Rabbit |
| Cyclin A1 | Cell Signaling | #4656 | 1:1000 | Mouse |
| Cyclin B1 | Cell Signaling | #12231 | 1:1000 | Rabbit |
| Cyclin E2 | Cell Signaling | #4132 | 1:1000 | Rabbit |
| γH2AX | Sigma-Aldrich | # 05-636 | 1:2000 | Mouse |
| GFAP | Cell Signaling | # 80788 | 1:200 | Rabbit |

**Supplementary Table 2.**
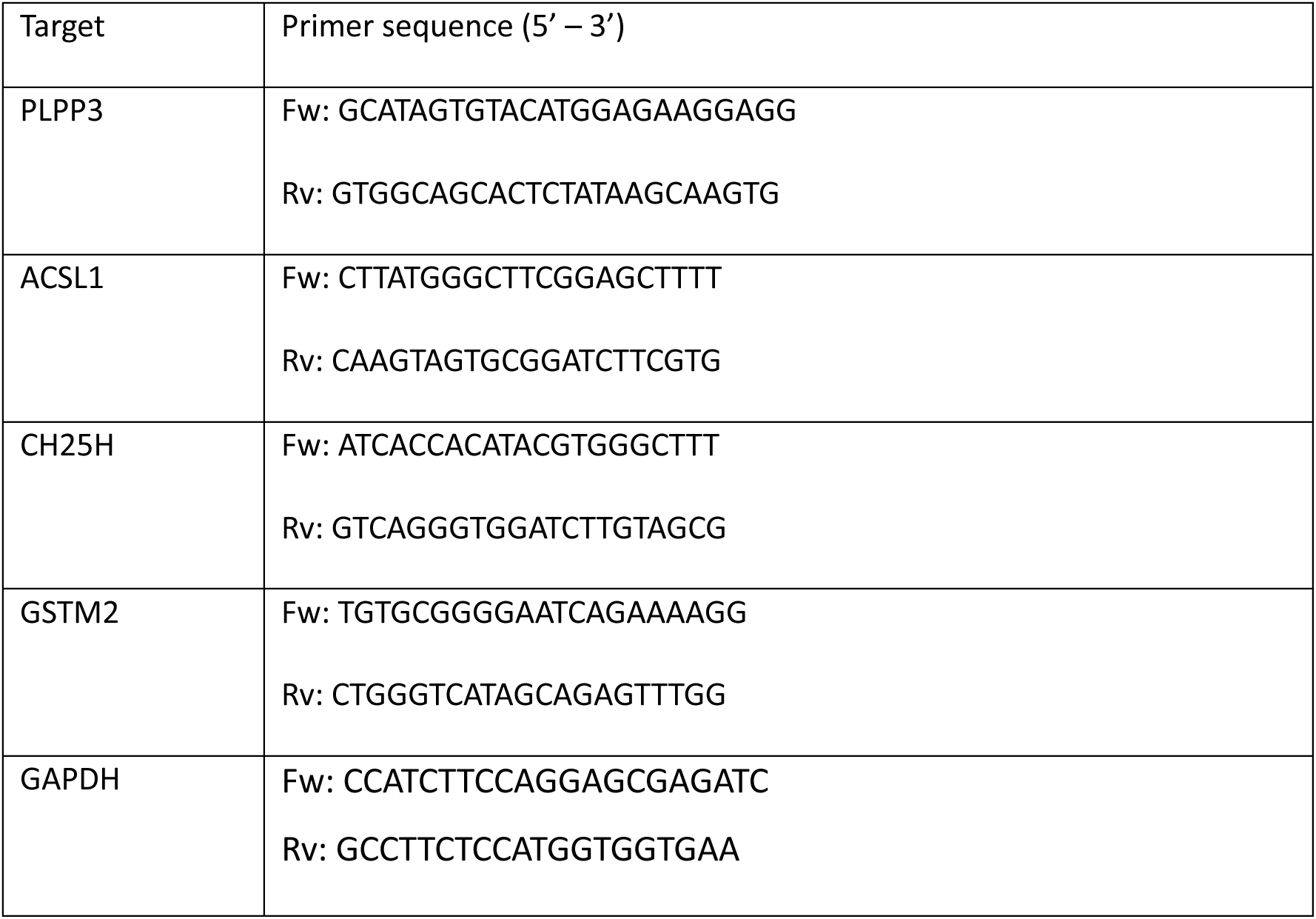
List of qPCR primers used in this study, indicating target gene and primer sequences.

